# Integrating Non-Spiking Interneurons in Spiking Neural Networks

**DOI:** 10.1101/2020.08.13.249375

**Authors:** Beck Strohmer, Rasmus Karnøe Stagsted, Poramate Manoonpong, Leon Bonde Larsen

**Affiliations:** SDU Biorobotics, Maersk McKinney Moller Institute, University of Southern Denmark

## Abstract

Researchers working with neural networks have historically focused on either non-spiking neurons tractable for running on computers or more biologically plausible spiking neurons typically requiring special hardware. However, in nature homogeneous networks of neurons do not exist. Instead, spiking and non-spiking neurons cooperate, each bringing a different set of advantages. A well researched biological example of such a mixed network is the sensorimotor pathway, responsible for mapping sensory inputs to behavioral changes. This pathway is also well researched in robotics where it is applied to achieve closed-loop operation of legged robots by adapting amplitude, frequency, and phase of the motor output. In this paper we investigate how spiking and non-spiking neurons can be combined to create a sensorimotor neuron pathway capable of shaping network output based on analog input. We propose sub-threshold operation of an existing spiking neuron model to create a non-spiking neuron able to interpret analog information and communicate with spiking neurons. The validity of this methodology is confirmed through a simulation of a closed-loop amplitude regulating network. Additionally, we show that non-spiking neurons can effectively manipulate post-synaptic spiking neurons in an event-based architecture. The ability to work with mixed networks provides an opportunity for researchers to investigate new network architectures for adaptive controllers, potentially improving locomotion strategies of legged robots.

## 1 Introduction

Current research employing neural networks for locomotion control tends to focus on homogeneous networks of neurons communicating through either graded signals (Aoi et al., 2017) or action potentials (Bing et al., 2018). However, studies indicate that biological neural networks utilize both communication strategies (Burrows, 1996) to achieve effective locomotion. Based on this, our research introduces a biologically-inspired non-spiking interneuron (NSI) model into a spiking neural network (SNN) to further increase biological fidelity. In nature, sensor neurons receive information from the external environment and pass it onto NSIs through current injections (Bidaye et al., 2018). This data is sent onwards by the NSI, affecting the membrane potential of the connected neurons through a graded signal (Burrows and Siegler, 1978). However, NSIs are not only translational units. They are also found to be the primary neuronal type in some animals such as the C. Elegans where communication through graded potentials is the main transmission method (Schafer, 2016). Thereby, interneurons are computational units in and of themselves. Figure 1 illustrates a simplified neural pathway depicting an analog input from the environment producing movement by an insect (A) and the equivalent pathway implemented in this study (B).

**Figure 1:**
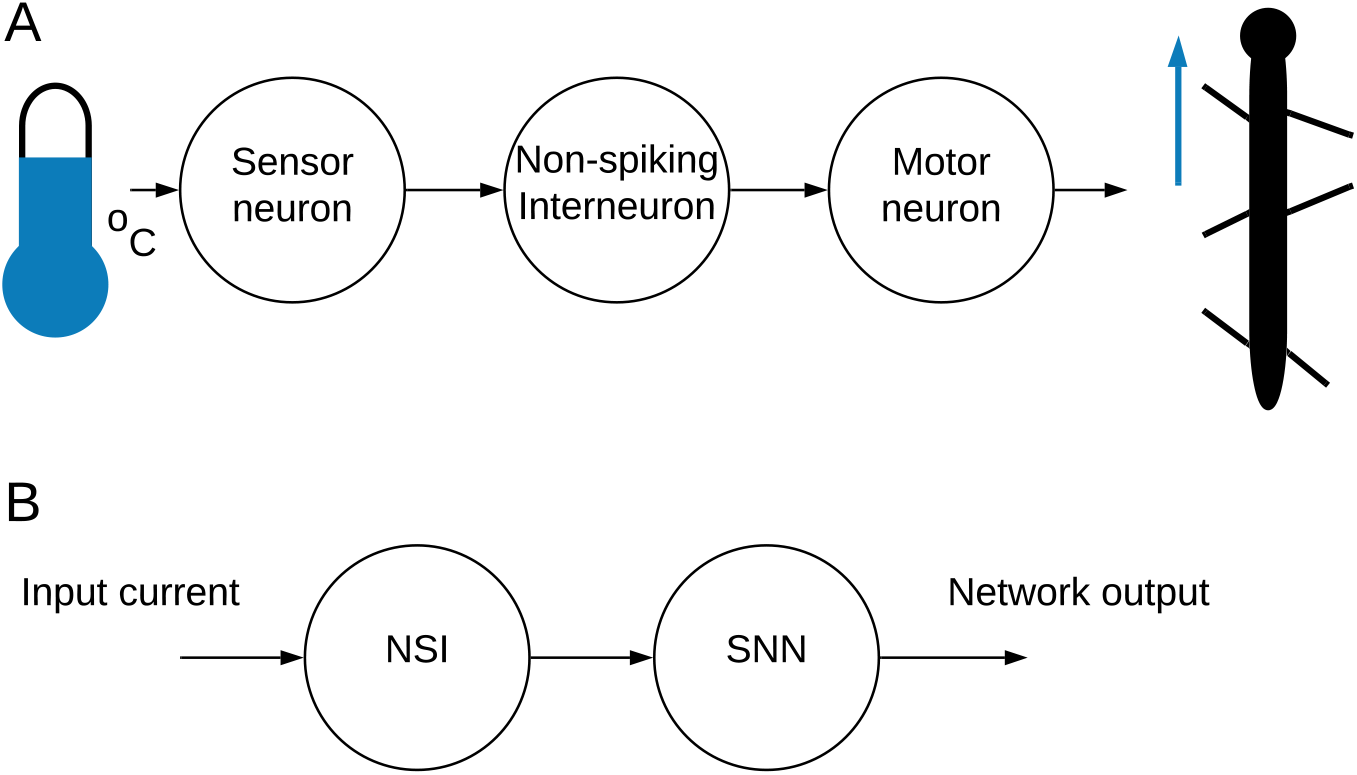
A: An illustration of a simplified sensorimotor neuron pathway. A signal is received from the outside world and sent onwards from the sensor neuron to the NSI through current injection. The NSI passes the information onto the motor neuron to generate movement. B: A high-level overview of the mixed neural network combining an NSI and SNN.

Individual neurons can be described using different models which try to capture the dynamics of a biological neuron. In this study, we use two spiking neuron models, one works in its intended fashion but the other operates within its sub-threshold range so that the membrane potential never surpasses the spiking threshold. Spiking neuron models try to replicate biological neurons by calculating the membrane potential of the neuron at each time step, this potential is affected by incoming spikes, bias current, and other parameters depending on the model’s equation. Classical non-spiking neuron models used in artificial neural networks (ANNs) attempt to replicate neuron dynamics using a transfer function such as the sigmoid function. These non-spiking models are able to map values but they cannot integrate an input over time without a recurrent connection. Therefore, our paper uses a spiking neuron model in a non-spiking “mode” to create the NSI.

SNNs typically communicate through action potentials commonly called spikes. Spikes allow information to be encoded through the frequency of spikes as well as the timing of individual spikes (Bohte, 2004). This creates the possibility to transfer more information within a spike train as compared to traditional ANNs (Bing et al., 2018). SNNs are used in our research so future work incorporating data-rich sensory input can take advantage of temporal features for encoding input information.

As SNNs are typically homogeneous, all communication is handled by passing spikes between neurons. If an analog sensor is added to the network, the information must be encoded into spikes for the network to understand. This is also true for an output signal which must be converted from spikes to continuous values to control a motor. The known encoding mechanisms found in nature include individual spike rate (Adrian, 1926), population activity (Panzeri et al., 2015), and precise spike timing (Bohte, 2004). Neural network engineers have applied each of these methods for translating sensory input, grouping the different approaches into three main categories: rate, population, and temporal coding respectively.

Each encoding category has recognized strengths. Population coding is able to relay more information than individual neurons so it can be useful for data-rich inputs (Mallot, 2013). On the other hand, temporal coding is particularly applicable for encoding streaming data as it quickly processes information (Petro et al., 2019) while maximizing the amount of information contained within the compressed data (Sengupta and Kasabov, 2017). Finally, while other methods are able to encode more information, rate coding has been suggested to be the best tool for handling input with high firing rates (Azarfar et al., 2018). In our work, rate coding is used to filter the output of the network from spikes to a continuous motor signal and we introduce the NSI model as a hybrid encoding method able to directly translate input by itself or work together with rate, population, or temporal methods to increase the amount of information encoded.

### 1.1 Related Work

ANNs have been shown to effectively manipulate amplitude, frequency, and phase in legged robots to create adaptive controllers (Thor and Manoonpong, 2019, Pitchai et al., 2019, Nachstedt et al., 2013, Barikhan et al., 2014, Schilling et al., 2013, Dürr et al., 2019). Thor and Manoonpong (2019) used an error signal to update synaptic weights for adaptation of frequency to optimize walking, resulting in increased efficiency and reduced tracking error. Pitchai et al. (2019) also created an energy-efficient control mechanism for a legged robot by using an ANN to shape network outputs coupled with a non-spiking central pattern generator (nCPG) to change frequency. Nachstedt et al. (2013) were able to create a self-tuning network using adaptive oscillators which allowed a robot to navigate a more complex environment. Barikhan et al. (2014) showed that a decoupled nCPG network using sensory feedback to adapt to the environment was able to handle changes in robot morphology and could coordinate movement with another robot when working on cooperative tasks. Schilling et al. (2013) used rules for coordination to adapt walking according to sensory input. Similarly, Dürr et al. (2019) developed an ANN to control a hexapod robot which relied on feedback for posture and rules for coordination. Their network produced emergent gaits, adjusting based on the posturing of the robot. The addition of non-spiking interneurons to ANNs has been investigated by Szczecinski et al. (2015). They reported control of a hexapod robot using interneurons modeled as classical non-spiking neurons to trigger different bio-inspired reflexes. The interneurons were used to control output oscillations, indicating that shaping nCPG outputs via interneurons to reproduce biological behaviors is possible. Our paper combines the use of NSIs and spiking neurons to update amplitude, frequency, and phase, as a step towards creating a more biologically plausible adaptive controller capable of interpreting temporal data.

Pure SNNs are also capable of manipulating output amplitude, frequency, and phase by updating different synaptic and neuronal characteristics (Strohmer et al., 2020). It was found that frequency could be changed by updating the value of the voltage threshold potential of the spiking central pattern generator (sCPG) neuron populations while amplitude was increased or decreased using the weight of the synaptic conductance to the motor neuron population (MNP). Finally, phase was determined by the network architecture and synaptic delays. However, as input currents to NSIs are known to change firing rates of connected motor neurons, reset rhythmic output (Bidaye et al., 2018), and update amplitude (von Uckermann and Büschges, 2009), it is useful to look into how they interact with an sCPG network to shape these outputs.

Woźniak et al. (2020) integrated spiking neurons into an ANN to take advantage of their power-saving potential and temporal data encoding capabilities. They implemented the spiking neurons as a “spiking neural unit” consisting of two non-spiking neurons, one of which handles integration of the membrane potential and the other to emit a spike. The spiking neural unit dynamics were modeled after the leaky integrate-and-fire model of spiking neurons. Their integration of the membrane potential was handled through a recurrent connection while spiking was mimicked using a step function. This approach differs from our implementation as it defined the spiking neurons as a combination of non-spiking neurons whereas we take the dynamics of a spiking neuron model and use it in the sub-threshold region to create a non-spiking neuron model.

Patil et al. (2015) took a similar approach to our research and created a non-spiking neuron modeled as a spiking neuron that communicated via graded potentials. Their work was based on the neural architecture of the C. Elegans which is mostly composed of non-spiking neurons though recent research indicates more advanced sensory systems in the worm might use spiking neurons (Liu et al., 2018). The paper showed that a mixed neural network could be built to mimic the escape response of the C. Elegans upon being touched externally. Their mixed network was implemented as neuromorphic hardware, simulating both non-spiking and spiking neurons using analog circuitry, as opposed to our research using simulation software to mathematically model neurons.

Our paper presents a novel encoding mechanism, sub-threshold encoding, that uses NSIs to relate values from analog sensory inputs to an SNN. Our proposed method is referred to as sub-threshold encoding because the output is a consequence of membrane potential fluctuations below the spiking threshold. In contrast to using current injections to directly manipulate network output, the momentary membrane potential of the NSI plays a role in how the post-synaptic neurons are affected. The main contribution of this research is the introduction and investigation into how these NSIs can be integrated into existing SNNs to shape network output.

## 2 Methods

Biological research indicates that sensory input to NSIs affects motor output (Büschges and Wolf, 1995). Furthermore, depolarizing currents received by NSIs are shown to reset biological central pattern generator (bCPG) rhythms (Bidaye et al., 2018). We can replicate these behaviors by creating an equivalent neural network representing the sensorimotor neuron pathway. In the neural network, the sensor neuron is replaced by an input bias current to the NSI (see Figure 1B) so that a change in input current represents a change in sensory information from the external environment. The NSI then relates this information to the connected SNN in order to adjust the network output. Figure 2B shows the block diagram of the selected SNN, an sCPG network. The architecture of a pair of mutually inhibitory neuron populations Is based on the biology of spiking oscillators driving an antagonistic muscle pair (Bidaye et al., 2017). The output from the sCPG network is spikes but an analog signal is required for control of a motor. Therefore, Figure 2A demonstrates how these spike events are converted to an analog signal using rate coding. This mimics the low-pass filtering performed by biological muscles (Hooper et al., 2007) by counting the amount of spikes occurring within a time window to produce an analog value. Figure 2C highlights the characteristics of the rate-coded output to be adjusted by the input to the NSI.

**Figure 2:**
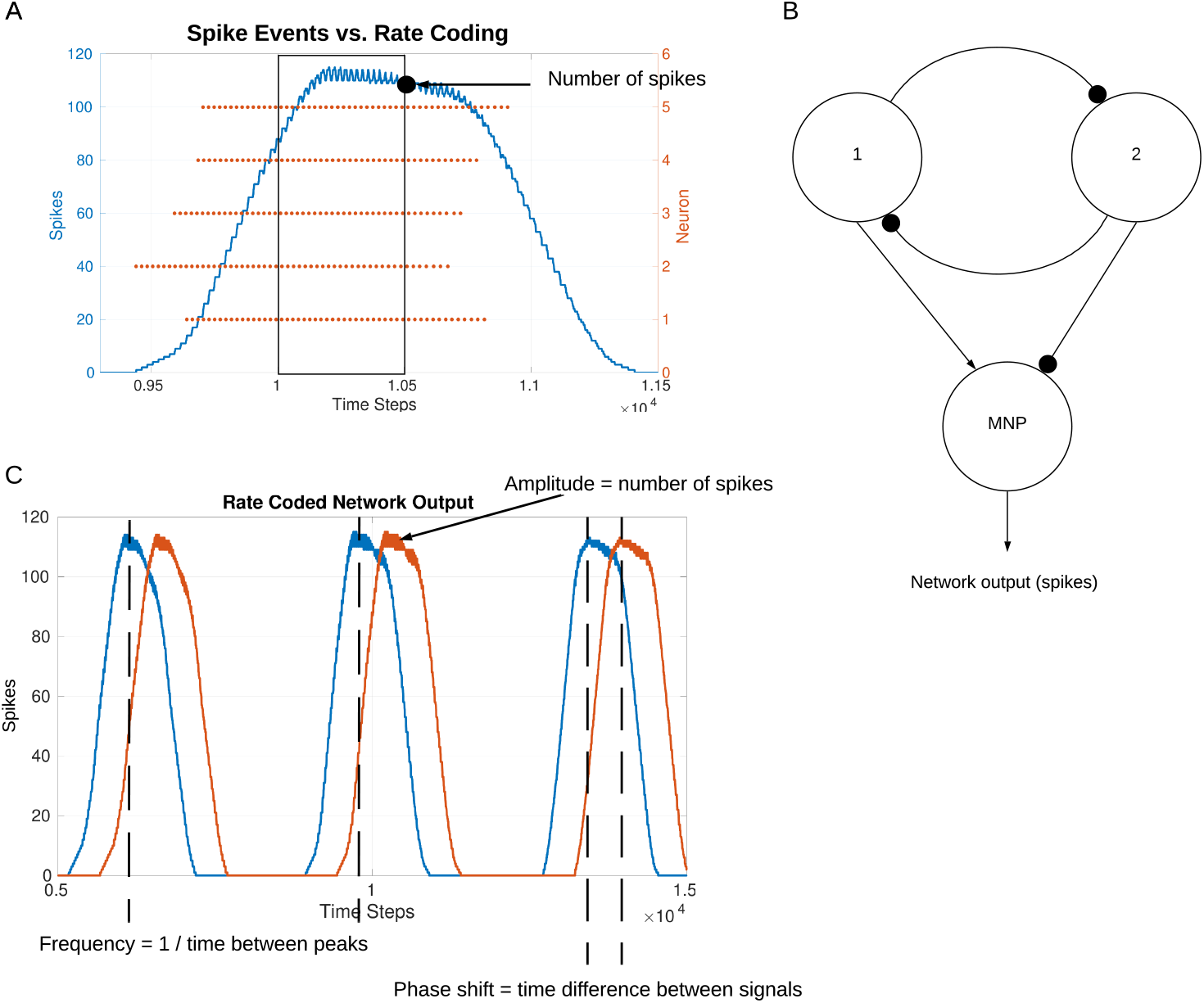
A: Illustration of how spikes are converted to analog values. The spike events for all neurons within a time window are counted. This value is used as the analog output value of the network. B: Block diagram of the sCPG network. Neuron populations 1 and 2 create the spiking oscillator, sending spikes to the MNP. The output spikes from the MNP are considered the network output. C: Illustration of network output characteristics, visualizing the terms amplitude, frequency, and phase.

A diagram of the implemented network is shown in Figure 3A. The network consists of an NSI capable of injecting current and manipulating the voltage characteristics of the post-synaptic neurons.

**Figure 3:**
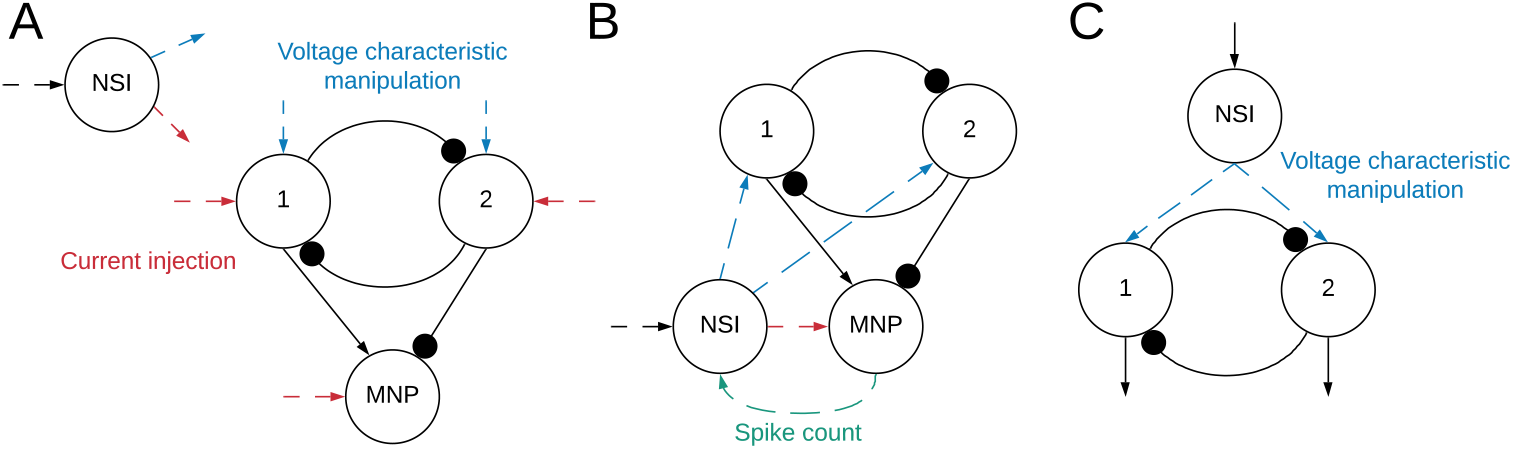
A: Block diagram outlining the network structure. Spiking neuron populations “1” (excitatory neu-ron population) and “2” (inhibitory neuron population) comprise the sCPG. The MNP is also spiking, it is regulated by an excitatory and inhibitory connection from the sCPG populations. The combination of the sCPG populations and the MNP create the complete sCPG network. The interneuron is non-spiking, communicating with the sCPG neuron populations through current injection and voltage characteristic manipulation. The MNP only receives current injections from the NSI. The neural network is altogether composed of 16 neurons, 5 neurons per population plus a single NSI. Red dashed lines indicate input and output of current between different neurons while blue dashed lines mark voltage characteristic manipulations. The dashed lines do not show direct connections because testing combined different configurations of current injection and voltage characteristic manipulation. B: Network for regulating output signal amplitude using the MNP spike count. The total number of spikes output by the motor neurons during the latest time step is sent to the NSI to determine the size of the current injection to the MNP. C: The event-based network consisting of an sCPG of two single AdEx neurons. The *V_th_* of each AdEx neuron is manipulated by an NSI based on spiking input.

The sCPG populations and MNP consist of 5 neurons each in order to produce a smooth, stable output with a minimum amount of neurons (Strohmer et al., 2020). The NSI is a single neuron so that all post-synaptic neurons connected to it receive the same inputs. This reduces the complexity of the experiments so that the tests focus on the communication from the NSI. The sCPG populations and MNP are comprised of adaptive exponential integrate-and-fire neurons to allow for bursting behaviors (Brette and Gerstner, 2005). Neuronal parameters are set based on regular bursting as outlined in Naud et al. (2008) and their dynamics are shown in equations (1) and (2).

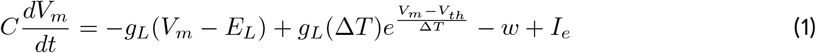

when *V_m_* > 0*mV* then *V_m_* → *V_reset_*

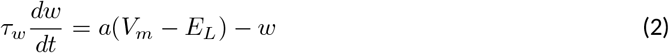

when *V_m_* > 0*mV* then *w* → *w* + *b*

where C is the membrane capacitance, *V_m_* is the membrane voltage, *E_L_* is the resting potential, *g_L_* is the leakage conductance, *I_e_* is the bias current, *a* is the sub-threshold adaptation conductance, *b* is the spike-triggered adaptation, Δ_*T*_ is the sharpness factor, *τ_w_* is the adaptation time constant, *V_th_* is the voltage threshold potential, *V_reset_* is the reset potential, and *w* is the spike adaptation current (Naud et al., 2008). Equation (1) defines the change in membrane potential per time step whereas Equation (2) outlines the current adaptation.

The NSI is simulated as a leaky integrate-and-fire neuron, the dynamics of the neuron are shown in Equation (3). The spiking threshold is set high enough to avoid spiking so there is no reset condition. The simpler neuron model is used for the NSI because the only necessary behavior is the integration of input current and leakage.

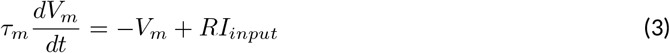

where *τ_m_* = *RC* is the membrane time constant, *V_m_* is membrane voltage, *I_input_* is input bias current, and *R* is membrane resistance.

Yang et al. (2013) show that the magnitude of the NSI output is a graded function of the difference between the interneuron’s membrane potential and its rest potential. Additionally, they find a linear correlation between the signal produced by the NSI and the response from the post-synaptic neuron. Based on this knowledge, we construct a relation between the NSI membrane potential and the effect on the post-synaptic neuron (Equations (4) and (5)).

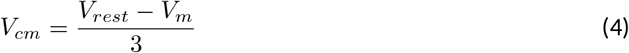

where *V_cm_* is voltage characteristic manipulation, *V_rest_* is rest potential, and *V_m_* is membrane voltage.

Equation (4) provides the offset to the voltage characteristic being adjusted for an sCPG neuron population. *V_rest_* is set to −60*mV* to stay consistent with biological findings (Graubard, 1978). The practical implementation of *V_rest_* uses the starting value of the NSI membrane potential as it fluctuates around the desired resting potential due to noise added to the system. The divisor in Equation (4) limits the voltage characteristic offset within a stable range. It is determined by dividing the biologically plausible 15*mV* NSI membrane potential fluctuation range (Burrows and Siegler, 1978) with the 5*mV* voltage characteristic manipulation range known to be stable for the sCPG network (Strohmer et al., 2020).

Equation (5) calculates the amount of current (*I_injection_*), in pA, to be added to the original current bias of the post-synaptic neuron.

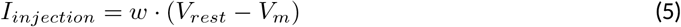

where *I_injection_* is current injection, w is synaptic conductance weight, *V_rest_* is rest potential, and *V_m_* is membrane voltage.

The synaptic conductance weight, *w*, scales the current injection from the NSI to the post-synaptic neuron. Conductance is measured in Siemens (S), the inverse of Ohms (*Ω*^−1^).

### 2.1 Time-Driven Experimentation

Two main categories of tests are performed on the network, excitatory and inhibitory. The input bias current (*I_input_*) to the NSI is always positive, this is the input current defined in Equation (3). However, the injection current (*I_injection_*) from the NSI to the post-synaptic neurons changes sign depending on test type, sending a positive current if excitatory and negative current if inhibitory. The value of *I_injection_* is determined by Equation (5). Testing is further broken into interactions between the NSI and post-synaptic neuron populations. The overview of these tests is outlined in Table 1 and visualized in Figure 4.

**Figure 4:**
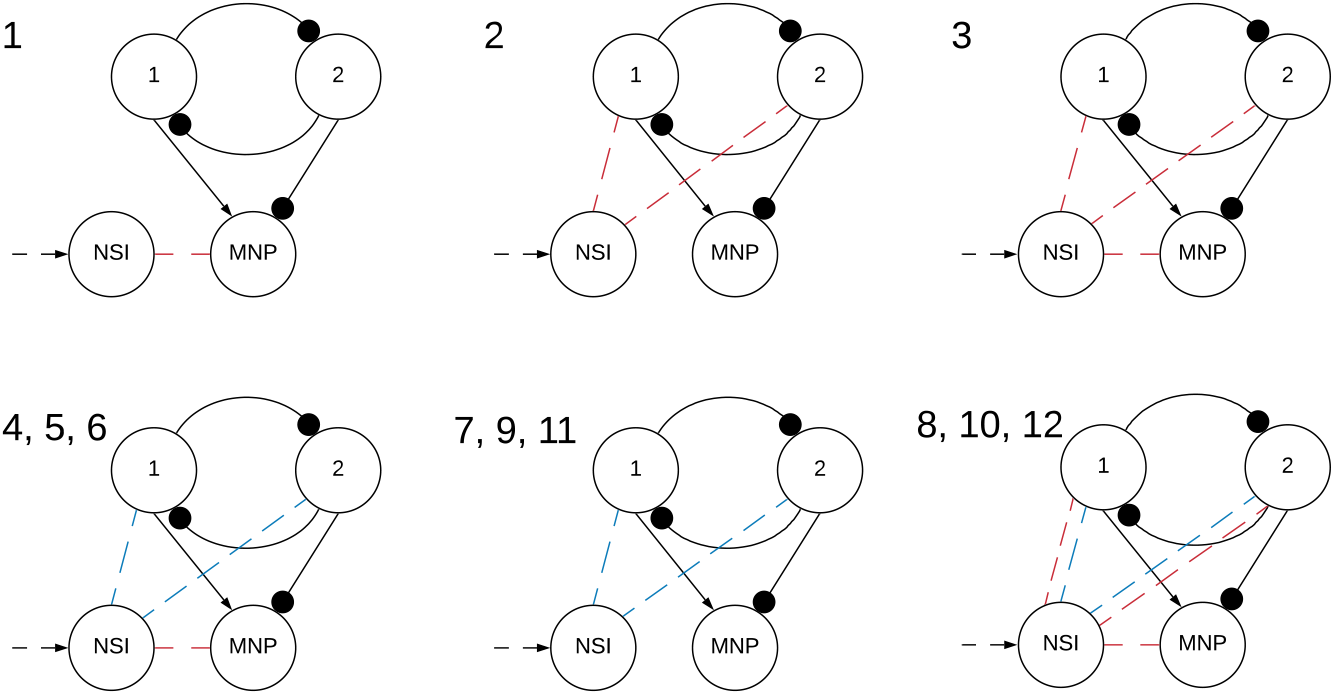
Visualization of NSI communication testing outlined in Table 1. Red indicates a current injection and blue indicates a voltage characteristic manipulation. Input bias current to the NSI is always excitatory (black dashed arrow). The numbers indicate the main test number corresponding with the depicted setup.

**Table 1:**
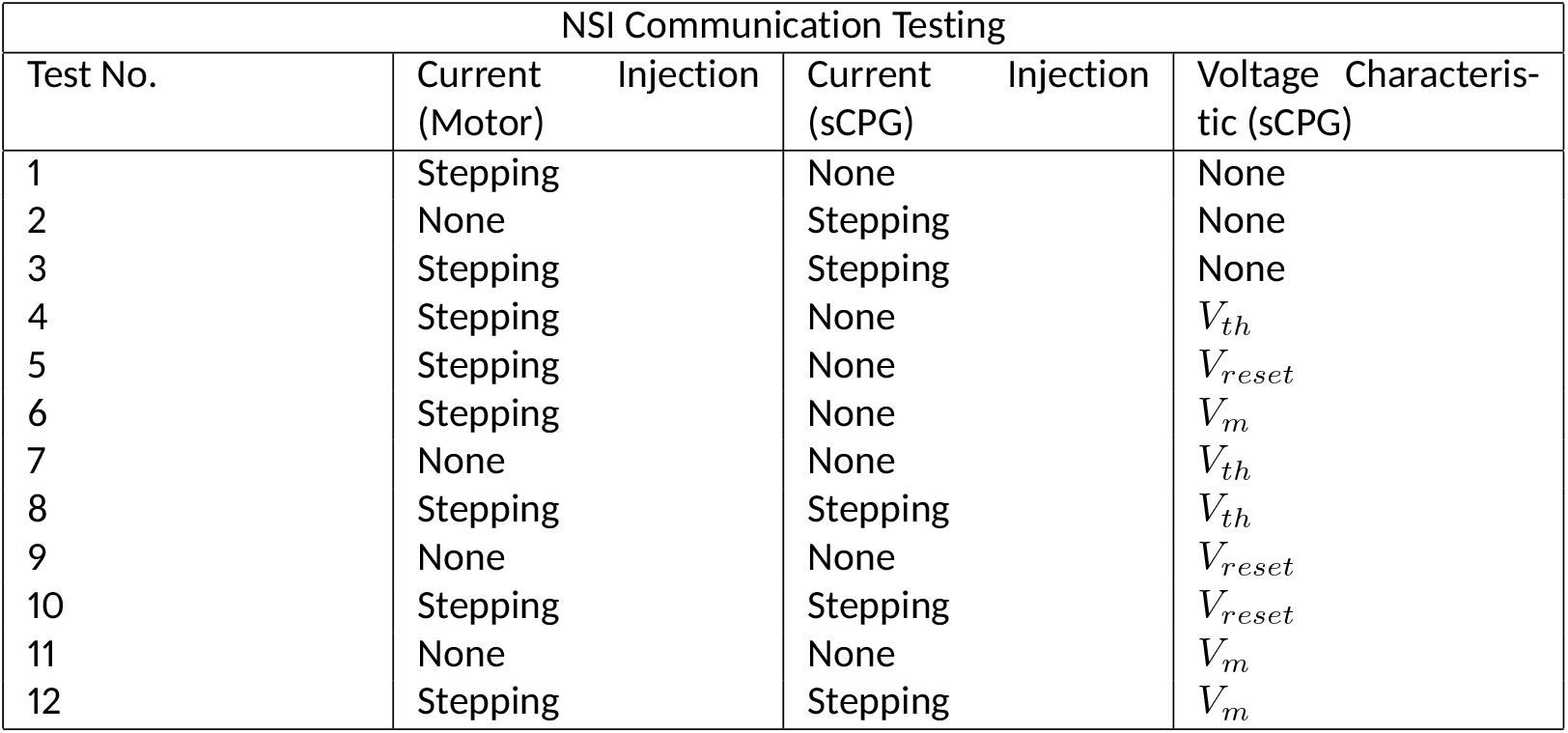
Test code reference

The tests step through different configurations of injecting current and manipulating voltage characteristics of post-synaptic neuron populations as seen in Figure 4. The voltage characteristics tests separately investigate adjusting voltage threshold potential (*V_th_*), voltage reset (*V_reset_*), and membrane potential (*V_m_*) within Equations (1) and (2). The amount of the offset is determined by the voltage characteristic manipulation Equation (4).

Each of the voltage characteristic manipulations is tested on individual sCPG neuron populations as well as both simultaneously, these are defined by test subcategories seen in Table 2 and visualized in Figure 5. When a test is running that does not update a particular voltage characteristic, the following values are used: *V_th_* = −51*mV* and *V_reset_* = −46*mV*. These values are known to produce a stable, regular bursting pattern by the sCPG network (Strohmer et al., 2020). The MNP never changes voltage characteristics, it only receives current injections because it is not involved in generating rhythmic patterns, only shaping the output.

**Table 2:**
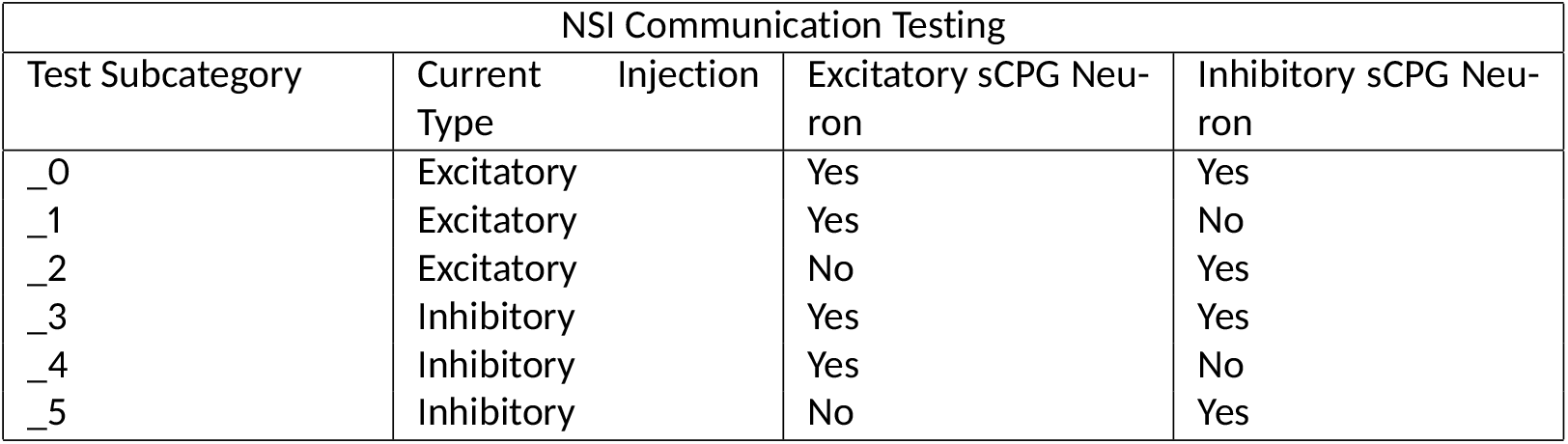
Test subcategory code reference

**Figure 5:**
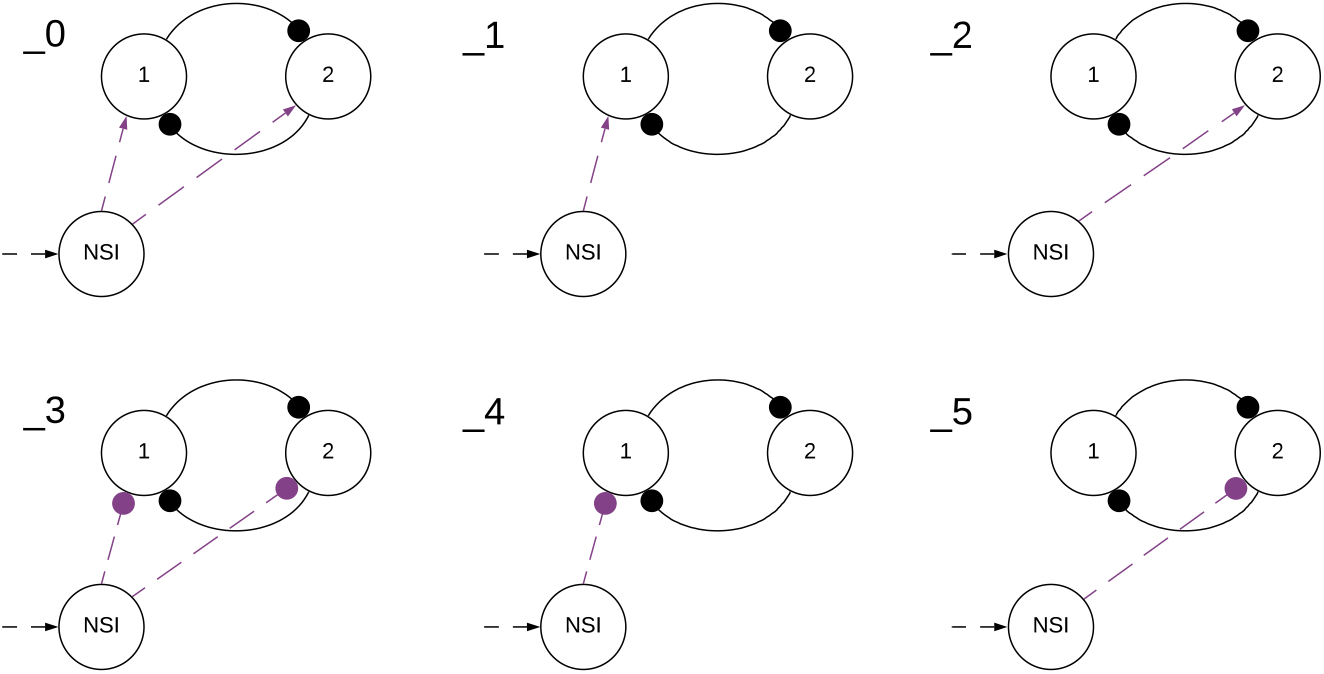
Visualization of NSI communication testing subcategories outlined in Table 2. Arrows are excitatory and solid circles are inhibitory connections. The connection may be a current injection, a voltage characteristic manipulation, or both depending on the main test number found in Table 1. This visualization shows how the subcategories determine the input to the sCPG populations. The numbers indicate the test subcategory corresponding with the depicted setup.

The test results are compared by plotting the rate-coded output of the MNP. A sliding time window of 5*ms* is used, counting all spikes from the MNP occurring within each time window to produce an analog value. The output signal from the MNP is considered the output of the network because this would be the signal used to control a motor when testing on a physical robot.

Static *I_input_* tests are performed first to find the maximum value for the conductance weight, *w* (Equation (5)). These static tests are not outlined in the above tables as they are only for parameter tuning. In order to find *w*, *I_input_* is set to the maximum value determined by the change in the NSI membrane potential, restricting it to the biologically plausible range of 15*mV* (Burrows and Siegler, 1978). Then *w* is increased by increments of 10*nS* until the maximum weight is found. The maximum is accepted as the largest weight value to produce an ideal output signal, this is further discussed in Section 3. This value is set as the maximum conductance weight (*w_max_*) for both excitatory and inhibitory tests. After this is found, all test configurations update *I_input_* at regular intervals to confirm a change in analog input is able to manipulate network output online. These stepping input current trials all use *w_max_* to calculate *I_injection_* in Equation (5). When the test is excitatory, *I_input_* starts at 0*pA* and ends at the determined maximum, the opposite is true when the test is inhibitory. This allows the system to be “excited” by reducing inhibition so the same general behavior can be expected at the output.

Neural Simulation Tool (NEST) (Jordan et al., 2019) is used to simulate the network and record test results. The simulation is run for 6*s* for all trials. The input current to the NSI updates every 1 second, allowing the network to settle after initial transients before the input changes again. *I_input_* starts at 0*pA* and ends at the maximum, 148*pA*. Thus each step adds 29.6*pA* of input current. Gaussian white noise current is added to all neurons in the network. The standard deviation of the noise current to the NSI is set to 25*pA* so that the noise current is comparable to the current steps. When testing with larger values than this, the output no longer reliably produces the necessary offsets for the voltage characteristic manipulation. For all other neurons, the standard deviation is 50*pA*. The standard deviation of the noise currents are selected and have not been tuned. A bash script is used to run tests in a reliable manner. The bash script and the python test script are available on GitLab (Strohmer, 2020).

After performing all of the test combinations for frequency and amplitude manipulation as outlined in Tables 1 and 2, one further test is performed to see how output phase is affected by switching between frequencies. The frequency is either held constant for 6*s* or toggled between 4*Hz* and 8*Hz* updating every second for 6*s*. There is no current injection to the motor population for these tests in order to examine phase as affected solely by frequency. This test differs from previous phase manipulations of an sCPG network where phase was determined by synaptic delay (Strohmer et al., 2020). Instead of trying to create a specific phase shift, this trial only confirms that phase is affected by changing frequencies and does not attempt to control it.

Insects use proprioceptive sensor neurons to understand the position of their limbs in relation to their body. These internal feedback loops can help an insect engage resistance reflexes to maintain posture (Tuthill and Wilson, 2016). This knowledge is used to implement a simplified network simulating an internal feedback loop (illustrated in Figure 3B) as a proof of concept application. The rate-coded output from the MNP determines the strength of the synapse connecting the NSI and MNP. A number of trials are run to determine an appropriate scaling factor for excitatory and inhibitory weight calculations.

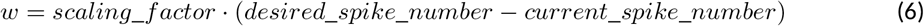

Varying the synaptic conductance weight, *w*, will regulate the output amplitude for this internal feedback loop. Equation (6) determines the *w* to be used in Equation (5) instead of using *w_max_*. The *scaling factor* is a value which allows for maximum effect without over-exciting or over-inhibiting the output. The *desired spike number* acts as a set point whereas the *current spike number* is the actual number of spikes from the MNP occurring in the last time step. In this network an inhibitory weight is applied if the number of spikes in the time window is above the desired number and an excitatory weight if below.

### 2.2 Event-Driven Experimentation

In order to confirm the NSI’s compatibility with a neuromorphic architecture, the network is also simulated on CloudBrain (Larsen et al., 2020). CloudBrain is a scalable event-based SNN simulation platform utilizing event stream processing technologies to communicate between neurons implemented as microservices on a cluster. The NSI in CloudBrain is implemented as an event-based leaky integrator neuron so that the neuron model is comparable to the NEST simulation. The neuron updates its voltage potential upon receipt of an input event according to the dynamics shown in Equation (7).

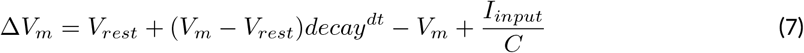

Where Δ*V_m_* is the change in membrane potential, *V_rest_* is the rest potential, *decay* is the decay factor simulating leakage, *dt* is the time since last incoming spike, *I_input_* is the input current from other neurons, and *C* is the membrane capacitance.

In addition to updating the voltage potential, the NSI saves the time stamp of last received input (*t_updated_*). Each microservice consists of a neuron and all incoming synapses so the NSI cannot directly access the voltage of the NSI. Instead, the NSI parameters (*V*, *V_rest_*, *V_m_*, *decay* and *t_updated_*) are sent to the synapse each time they are updated. Based on this information, the synapse asynchronously calculates the exact voltage potential of the neuron using Equation (8).

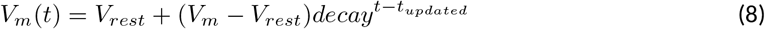

Where *t* is the current time, *V_m_* is the voltage potential, *V_rest_* is the rest potential, *decay* is the decay factor simulating the leakage, and *t_updated_* is the time at which the parameters were updated.

A network consisting of an NSI and an sCPG is created in CloudBrain to confirm the NSI is able to update the voltage threshold potential of the post-synaptic neurons in an event-based architecture. In this setup, the sCPG is comprised of two single neurons mutually inhibiting each other. Figure 3C shows the block diagram of the network in CloudBrain. The input to the NSI is spikes since it is an event-based architecture. The input spikes are the equivalent of the the stepping input current used in the time-based NEST simulation. A step function is encoded to spikes using Ben’s Spiker Algorithm (Schrauwen and Van Campenhout, 2003) for transmission to the NSI. In turn, the NSI updates the *V_th_* of the sCPG neurons based on the frequency of spiking input. The *V_th_* of the sCPG neurons is caculated using a linear mapping of the voltage potential. This allows us to control the upper and lower limits of the *V_th_*. The NSI’s *V_m_* is recorded for comparison with the *V_th_* and spiking output of the sCPG populations.

## 3 Results

Simulating with a constant *I_input_* of 148*pA* shows the NSI membrane potential starts close to the rest potential, −60*mV* and ends around −45*mV*, though these values vary slightly due to noise. This gives us the maximum allowable *I_input_* that restricts the NSI’s membrane potential within a 15*mV* range.

The maximum synaptic conductance weight for the NSI output synapse is determined to be *w_max_* = 70*nS*. The effect on amplitude as compared to a lower weight of 2*nS* is visible in Figure 6. Increasing *w* to 80*nS* produces unwanted lifting behavior from the excitatory current injection where the rate-coded output does not always return to zero. This result can be seen in Figure S1 in the supplementary material.

**Figure 6:**
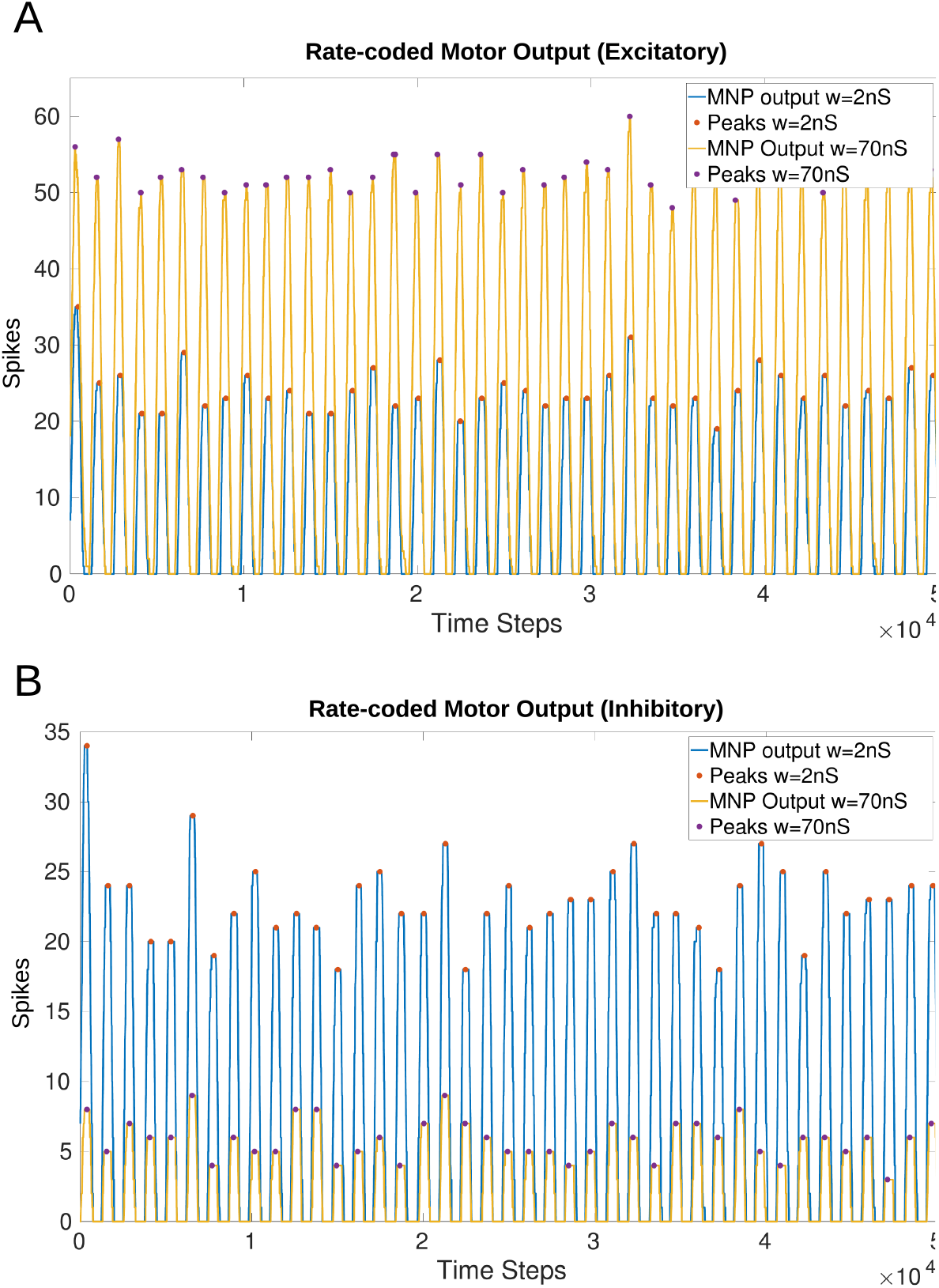
Comparison of the output from the MNP when *w* in Equation (5) is 2*nS* versus 70*nS* with a constant maximum *I_input_* of 148*pA* into the NSI (Equation (3)). There is a visible change in amplitude without affecting the frequency or phase.The peaks (local maxima) are marked with a filled circle. A: Injects excitatory current to the MNP; B: Injects inhibitory current to the MNP;

Figure 6 shows that amplitude is affected by a change in *I_injection_* without altering the frequency or phase of the output for both excitatory (A) and inhibitory (B) tests. The filled circles on the plots illustrate the peak values (local maxima) of the rate-coded output. The difference in average peak value when comparing 2*nS* versus 70*nS* is 28.37 spikes when excitatory and 16.69 when inhibitory.

Table 3 shows the progressive increase in average number of spikes per 5*ms* time window based on the size of *I_injection_* to the MNP. This reveals that the change in the number of output spikes due to *I_injection_* is scalable over a range.

**Table 3:** Average maximum number of spikes per time window when injecting excitatory or inhibitory current to the MNP. *I_injection_* shown is an approximation, calculated using Equation (5) where *V_m_* = −45*mV* and *w* is as defined in the table. *I_injection_* is positive when excitatory and negative when inhibitory. The average peak number of spikes increases and decreases with excitation and inhibition respectively.

| Calculated current injection, $I_{injection}$ (pA) | Conductance, $w$ (nS) | Average peak (excitatory) | Average peak (inhibitory) |
| --- | --- | --- | --- |
| 30 | 2 | 24.26 | 23.00 |
| 150 | 10 | 26.50 | 20.76 |
| 300 | 20 | 29.33 | 18.10 |
| 450 | 30 | 32.19 | 14.43 |
| 600 | 40 | 36.33 | 12.74 |
| 750 | 50 | 41.14 | 10.64 |
| 900 | 60 | 46.94 | 8.02 |
| 1050 | 70 | 52.63 | 6.31 |

### 3.1 Online Amplitude Manipulation

The online manipulation of *I_injection_* to the MNP results in a change in the number of spikes per time window for both excitatory and inhibitory tests (Figure 7A and B). The addition of *I_injection_* to both of the sCPG populations changes the behavior of spiking, shifting the phase in comparison to the static input current tests (Figure 7C, D, E, and F). Additionally, the strong inhibition of both sCPG neuron populations leads to complete suppression of the output.

**Figure 7:**
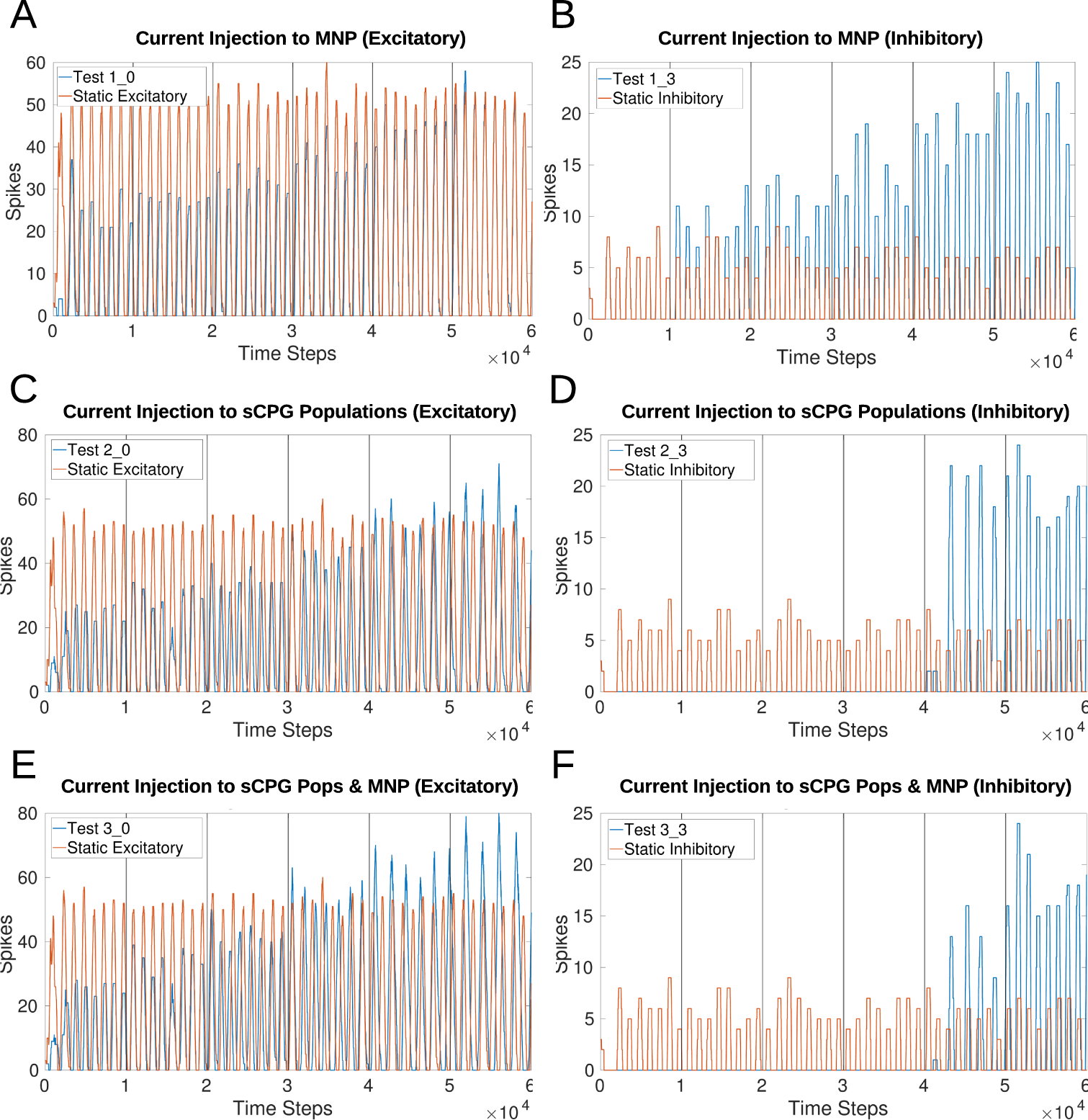
Comparison of various current injection configurations to the original static input test. For the stepping tests 1, 2 and 3, *I_input_* to the NSI increases every 1,000 time steps (1 second). Outputs show that current injection to the MNP changes the amplitude but the addition of current to the sCPG neurons creates a phase shift to the output. A and B: Baseline stepping current tests injecting current from the NSI to the MNP; C and D: *I_injection_* from the NSI to both sCPG populations; E and F: *I_injection_* from the NSI to both sCPG populations and the MNP; Excitatory tests are on the left while inhibitory tests are on the right.

Injecting current to one sCPG population at a time creates instability in the system. Figures S4A and B in the supplementary material show that the excitation of the excitatory sCPG population produces lifting behavior when *I_injection_* reaches a specific threshold. Likewise, an increase in excitation of the inhibitory neuron population dampens MNP spiking.

All figures in the supplementary material present both excitatory and inhibitory trials. Excitatory current injections produce a larger change in amplitude whereas inhibition suppresses output up to a certain point in many tests.

### 3.2 Online Frequency Manipulation

Figure 8 compares frequency manipulation of the output with and without current injection to the MNP. Updating either *V_th_* or *V_reset_* of the sCPG neuron populations points to a linear relationship with frequency.

**Figure 8:**
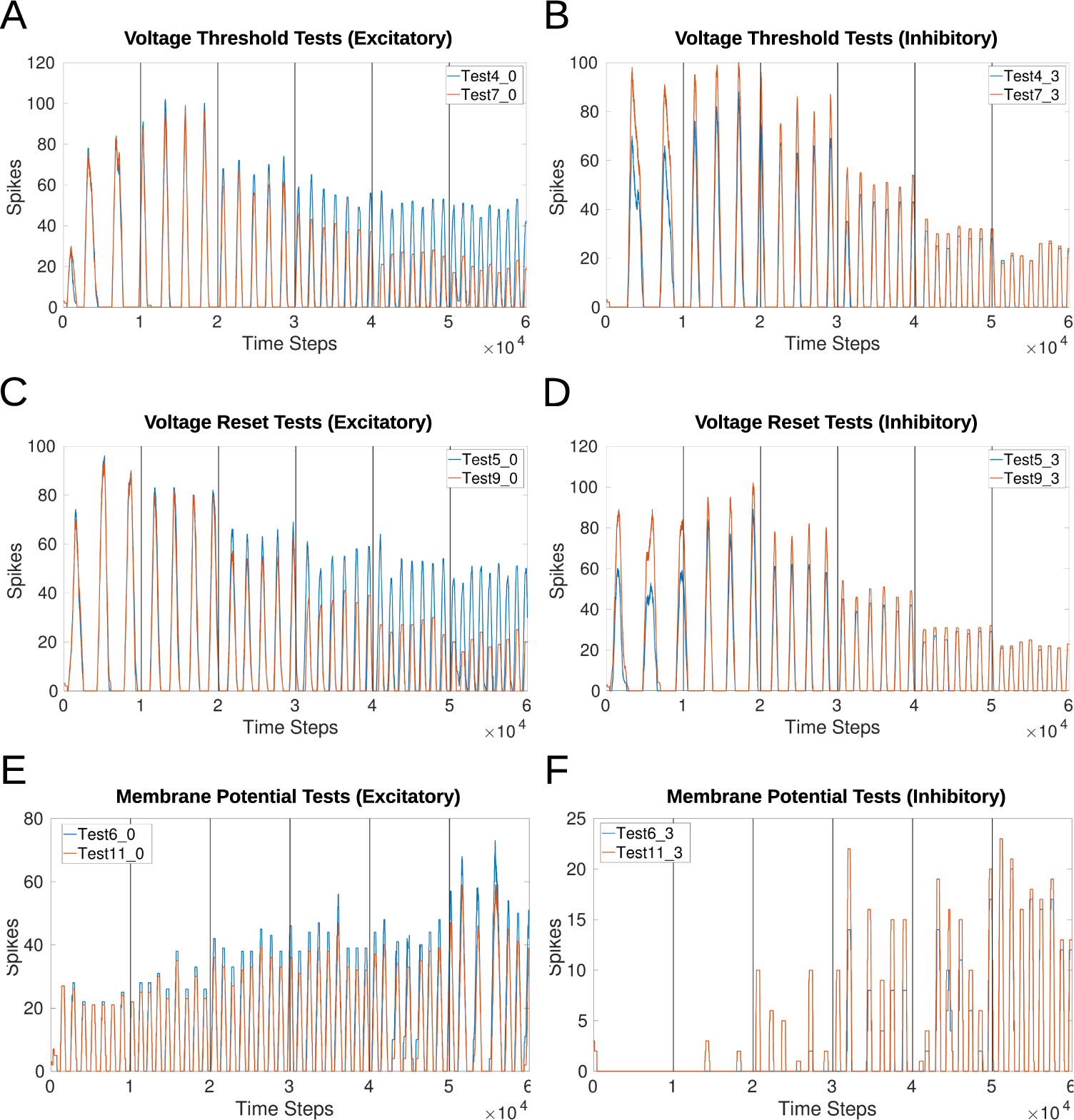
Comparison of voltage characteristic manipulation with and without current injection to the MNP. Network behavior remains the same when current is injected to the MNP during frequency changes for all voltage characteristic manipulations. Frequency changes linearly during *V_th_* and *V_reset_* tests. Excitatory and inhibitory tests are compared to their respective counterparts. Vertical lines indicate when a change in input current occurs once per second. A and B: Test 4 includes current injection to the MNP, Test 7 only updates voltage threshold potential; C and D: Test 5 includes current injection to the MNP, Test 9 only updates voltage reset; E and F: Test 6 includes current injection to the MNP, Test 11 only updates membrane potential.

Based on the voltage characteristic manipulation calculation (Equation (4)), *V_th_* and *V_reset_* are either increased or decreased by approximately 1 at each time step. Figures 8A, B, C, and D show that their manipulation results in an increase in frequency by the addition of approximately 1 peak per second, implying a linear relationship. Figures 9A, B, and C reinforce the expectation of a linear relationship between frequency and the change in *V_th_* or *V_reset_*. By contrast, adding an offset to *V_m_* does not reliably change the frequency and introduces instability as the current injection increases. Figure 9D shows that the frequency does not trend in a particular direction for the *V_m_* manipulation. Additionally, as the inhibitory test’s output is suppressed for much of the duration of the trial, there are fewer data points as compared to other tests.

**Figure 9:**
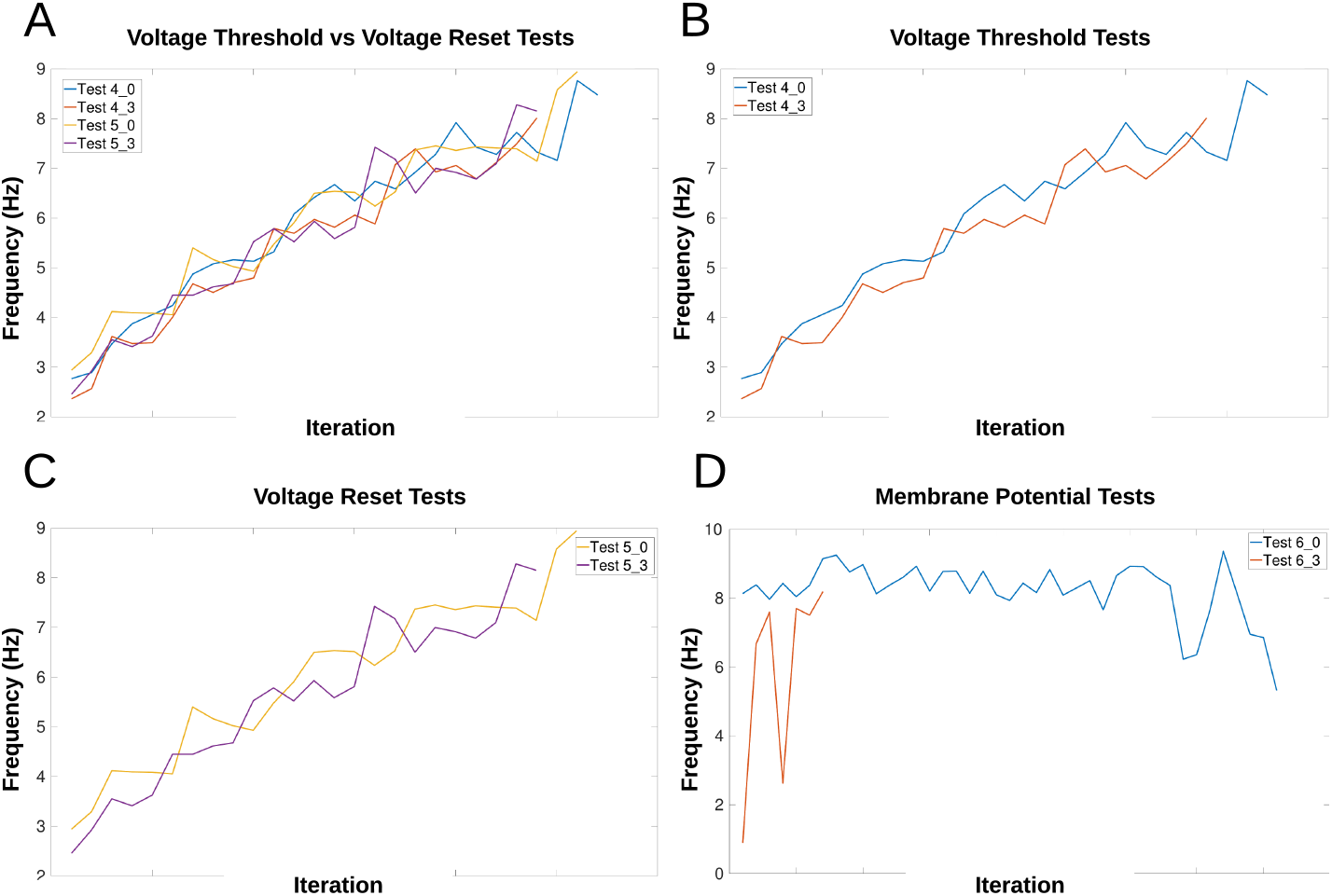
Calculation of frequency for each voltage characteristic manipulation using the time difference between peaks. Double peaks are removed as outliers. The x-axis only represents iteration through the array by subtracting the previous array element from the current element, it does not indicate time. The frequency of the MNP output trends upwards for A, B, and C, indicating that peaks occur more frequently when the absolute difference in voltage between *V*_*reset*_ and the *V*_*th*_ is decreased. The suffix _0 is used for excitatory tests and _3 for inhibitory tests. A: Compares peak frequencies of *V*_*th*_ and *V*_*reset*_; B: *V*_*th*_; C: *V*_*reset*_; D: *V*_*m*_;

Figure 9A highlights the similarity in the frequency change when comparing *V_th_* and *V_reset_*. This is also true whether the test is excitatory or inhibitory. Table S1 in the supplementary material shows the exact start and end frequency values for the voltage characteristic manipulation tests. The starting frequency for *V_th_* and *V_reset_* tests is between 2.36-2.94*Hz*, ending between 8.01-8.94*Hz*. This reveals frequency can be affected by a factor of 3.

Amplitude is also affected by frequency, the average number of maximum spikes for each frequency can be seen in Table S2 in the supplementary material. The lower the frequency, the more spikes are counted per time window, resulting in a higher value of the rate-coded output. The relationship does not appear to be linear as the number of spikes is reduced by a larger amount when comparing the lowest 3 frequencies as compared to the highest 3 frequencies.

### 3.3 Online Phase Manipulation

Phase is affected by frequency manipulation as seen in Figure 10. Figures 10A and B both show the initial 1,000 time steps of the simulation produce the exact same results for amplitude and phase. However, after a frequency change the phase shifts.

**Figure 10:**
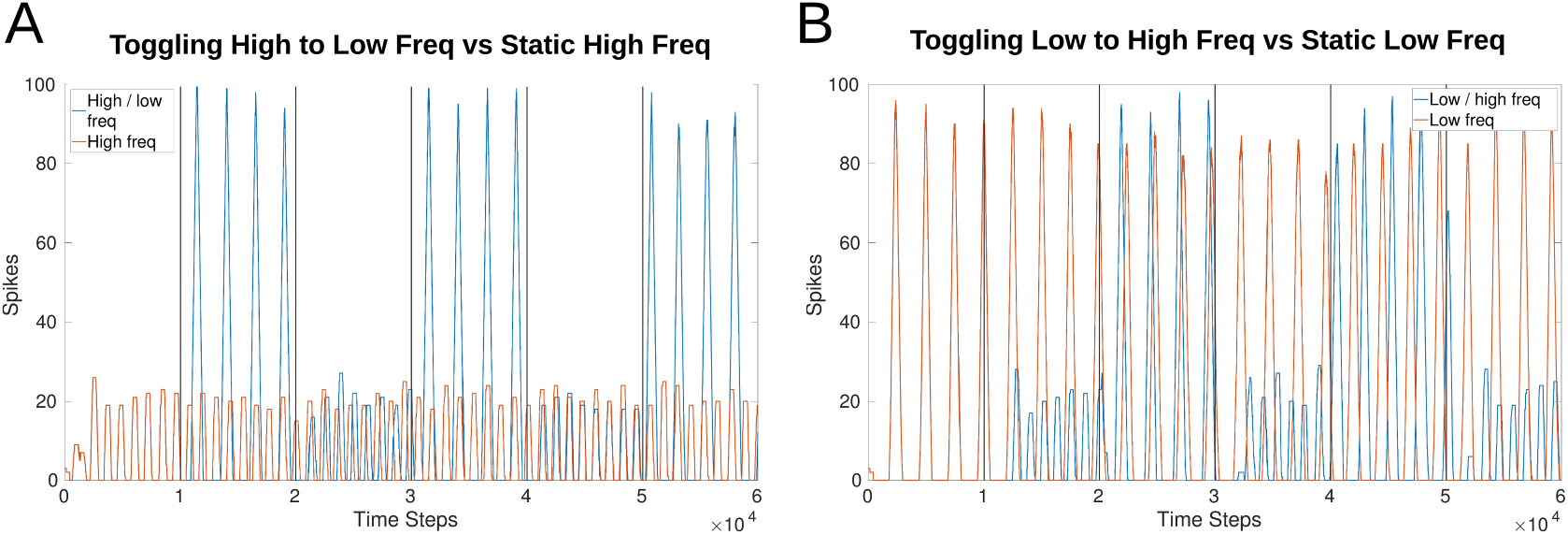
Comparison of output phase when toggling between frequencies versus a constant frequency. Both plots show the first 1,000 time steps (equivalent of 1 second) are exactly the same as the toggling (blue) and constant (red) frequency plots follow each other. After a change in frequency, the phase shifts so that they no longer follow each other exactly even though the frequency returns to the original speed. A: Compares a constant high frequency (red) to a toggling frequency (blue). B: Compares a constant low frequency (red) to a toggling frequency (blue).

### 3.4 Online Amplitude Regulation using Internal Feedback

Figure 11 shows the effects of amplitude regulation when updating the frequency of the network by adjusting *V_th_* in Equations (1) and (2). *V_th_* changes based on the offset determined by Equation (4). The scaling factors found to have the most effect without over-inhibition or over-excitation are 50 for inhibition and 2 for excitation (insert into Equation (6)).

**Figure 11:**
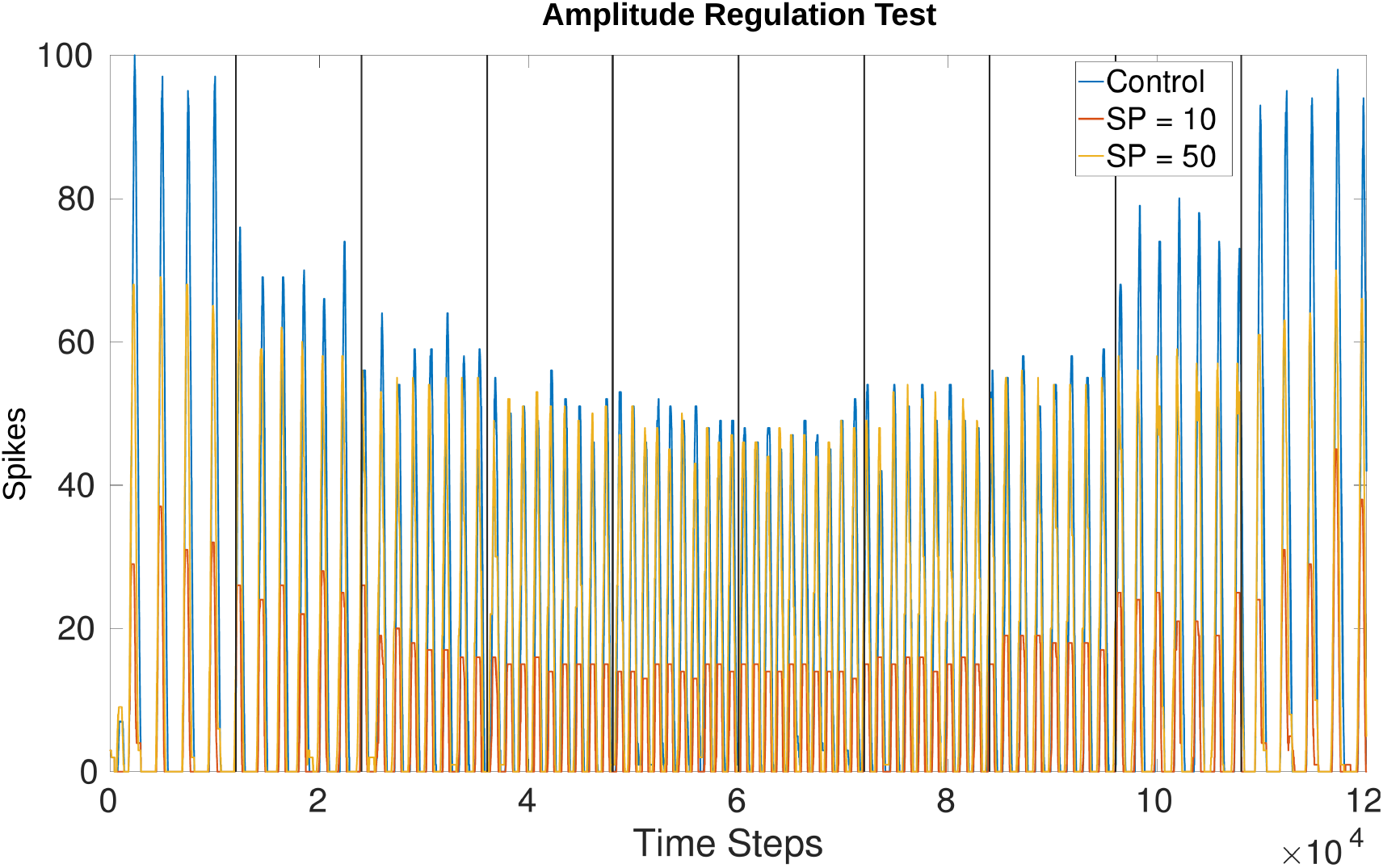
Comparison of amplitude when running the network unregulated versus creating a set point (SP) at 10 spikes and 50 spikes. The SP in this plot is equivalent to the “desired spike value” referred to in Equation (6). There is more overshoot for lower frequencies as compared to higher frequencies but the change in amplitude is visible from the control experiment where *w_max_* is applied uniformly. Vertical lines indicate when a change in voltage threshold potential occurs thereby changing the frequency. The threshold updates every 1,200 time steps (1.2*s*) creating a total of 10 divisions. *V_th_* starts at −54.81*mV* producing a frequency of 3.77*Hz* and increases to −50.79*mV* producing a frequency of 9.14*Hz*. After 6,000 time steps *V_th_* decreases again, returning to a value of −54.74*mV* producing a final frequency of 4.00*Hz*. The middle two trials from 4,800-7,200 time steps are held at the highest *V_th_*, around −50.79*mV*. The values fluctuate by hundredths of a *mV* because *V_th_* is affected by current noise.

The amplitude based on a “desired spike value” (see Equation (6)) of 10 versus 50 confirms that the network is monitoring the output and adjusting. This is seen more drastically when comparing the control experiment (originally test 4_0) and the trial with a desired spike value of 10. Regardless of whether the test is increasing or decreasing frequency, the amplitude is adjusted to a similar value. Figure 11 confirms the number of spikes per time dow is consistent based on the desired value and the frequency. Furthermore, the phase is unaffected by this amplitude regulation.

### 3.5 Event-Driven Manipulation

The step input to the NSI resulting in an increase in *V_th_* of the AdEx neuron is seen in Figure 12A. The y-axis is adjusted for each graph so the height of the curves should not be directly compared. However, the time axis is common to both graphs and shows the *V_th_* follows the change in *V_m_* almost immediately.

**Figure 12:**
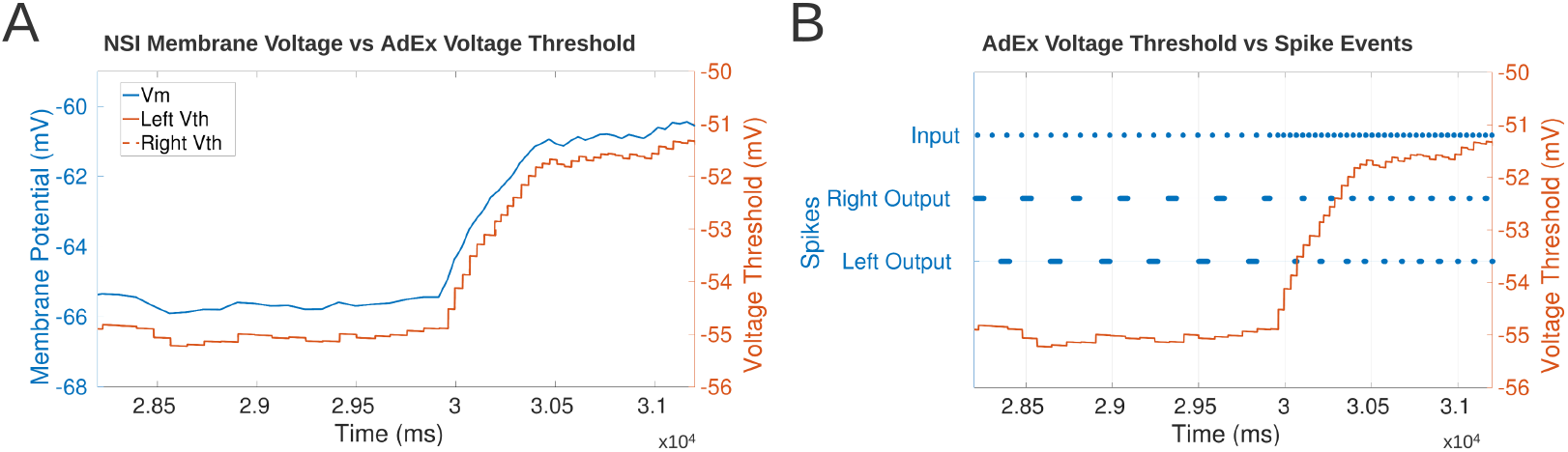
A: Comparison of the NSI’s *V_m_* to the post-synaptic AdEx neuron’s *V_th_*. The plot shows *V_th_* follows *V_m_* confirming that the neuronal characteristics are updated based on the membrane potential of the NSI. The AdEx neurons are defined as “left” and “right” based on their relative position in the sCPG. There is no noise on the synapses to the AdEx neurons so the *V_th_* of each neuron follows each other exactly thereby obscuring the view of the right neuron underneath the left. B: Comparison of each AdEx neuron’s *V_th_* to the input and output spike events. The spikes are not rate-coded because there is only one neuron per population. *V_th_* increases when the input spike events increase which leads to an increase in bursting frequency of the AdEx neuron. The left and right neurons spike 180 out of phase as expected based on the sCPG architecture of mutual inhibitory coupling.

Figure 12B confirms the output frequency of the AdEx neuron is increased when the input spike bursting frequency increases. The graph shows the increase in input bursting frequency starts ramping up the *V_m_* of the NSI. This promotes an increase in output bursting frequency from the AdEx neurons. The AdEx neurons are confirmed to spike out of phase with each other and increase to the same bursting frequency.

## 4 Discussion

The results confirm our model of an NSI is capable of shaping output and setting rhythmic patterns of an sCPG network based on a changing analog input. The amount of influence the NSI has on the output is constrained since the change in membrane potential must stay within the biologically plausible 15*mV* range. However, the range still allows at least a doubling of average output spikes per time window from the MNP and triple the frequency when moving from a low to high input current to the NSI. The output from the MNP also dictates usable parameter ranges. The lifting behavior observed when using a synaptic conductance larger than 70*nS* means that the output cannot always return to zero due to some neurons always spiking. This is not ideal as an output signal so the maximum conductance to the post-synaptic neurons is limited to 70*nS*.

The baseline stepping current tests (1_0 and 1_3) confirm amplitude can be manipulated online without changing the behavior of the system by updating the injection current to the MNP. Table 3 shows that the difference in amplitude can be controlled using synaptic conductance. It can logically be concluded that the average number of maximum spikes per time window when no current is injected is between 23 and 24.26. The number of spikes either increases or decreases from this starting point depending on if the connection is excitatory or inhibitory. The difference between maximum spikes over the range of current tested is steady, reinforcing the conclusion that amplitude is reliably altered by current injection. If the synaptic weight is changed in addition to the current injection, the average number of spikes per time window can be regulated from 6.31 to 52.63. This configuration increases the average spike difference to a factor of 8 (52.63/6.31 = 8.34).

Tests involving a current injection to the sCPG neuron populations produce a phase shift. This is expected because they are the populations driving the rhythmic output. The delay generated could be an exploitable feature when coordinating multiple joints. However, the exact effect has not been calculated and the delay changes based on the level of current injection (see Figure 7).

The linear relationship found between frequency and voltage threshold potential seen in Figure 9 is consistent with previous research by Strohmer et al. (2020). It follows that reset potential has a similar relationship to frequency as voltage threshold potential. Both characteristics change the amount a neuron’s voltage potential must change before reaching the spiking threshold. Adding an offset directly to a post-synaptic population’s membrane potential introduces discontinuities, creating small “jumps” based on the value of the offset. The frequency does not appear to be linearly correlated with the change in *V_m_* when looking at Figure 9D. However, this might be due to the limited range of the offset tested in this study.

The comparison of frequency to the maximum amount of spikes depends on the size of the time window chosen for rate coding as well as the number of neurons within the motor population. Likewise, a larger number of neurons in the motor population increases the maximum possible spikes. The results shown in this study are based on a time window of 5*ms* and a MNP size of 5. When comparing the start and end frequencies for Table S1 against the average frequencies in Table S2, the static tests show a higher starting, frequency (both tables are located in the supplementary material). The static tests average frequency over a total of 5 seconds, removing the first second in case of transients. However, Test 4_0 updates the frequency every 1 second so the minimum frequency and maximum frequency shown are not averaged but single calculations. The averaging process most likely accounts for this difference in start and end frequencies. This limits the reliable change in frequency to a difference of 2.8*x*.

Toggling between a low and high frequency over a single trial as compared to a constant frequency produces a phase shift as expected due to the alteration of the output signal’s period. This phase shift is with respect to the sCPG network’s own output from the MNP and does not indicate how frequency changes might affect a larger system with connected oscillators. However, these results imply that an NSI is able to reset rhythmic behavior of an sCPG network in accordance with biological research (Bidaye et al., 2018).

Trials that only affected a single sCPG neuron population at a time created network instability, most likely due to the change in network dynamics. The sCPG populations are mutually inhibitory so the excitation or inhibition of one population affects the balance of the network. This is particularly visible in Figure S4 in the supplementary material where over-excitation of the excitatory neuron causes lifting and over-inhibition of the excitatory neuron suppresses network output. Based on these results, a stable and predictable output requires updating both sCPG neuron populations with the same current injection or voltage characteristic manipulation. This conclusion can only be inferred for a mutually inhibitory sCPG architecture.

Reviewing the frequency tests with and without current injection to the motor population shows that the amplitude adjusts based on the level of current injection without affecting frequency. Furthermore, the frequency is equally affected by a change in *V_th_* or *V_reset_* but biological research shows evidence that voltage threshold adaptation is a strategy used by neurons to change firing rate (Azarfar et al., 2018). Therefore, based on research findings and biological research, our recommended approach for parameter manipulation of a mutually inhibitory sCPG network is to inject current to the MNP while updating the voltage threshold potential of the sCPG neuron populations.

The regulation of amplitude is a building block for developing closed-loop adaptive controllers with NSIs. The proof of concept network implemented shows that NSIs are capable of controlling output based on live feedback. The method is solely for demonstration purposes and is not necessarily biologically plausible though there is evidence that short-term plasticity affects rhythmic outputs (McDonnell and Graham, 2017) and that the synaptic weight influences the magnitude of the post-synaptic potential (Burrows, 1996). Testing the network reveals visible overshoot from the desired set point but this is expected because the spike value must surpass the set point before inhibition initiates. The lower frequencies are also worse at reducing the number of spikes, this is probably because of the significant excitation received from the sCPG populations. As can be seen in the control experiment, the maximum numbers of spikes per time window can reach 100 for the lowest frequency trial. It is more difficult to dampen the effects of these high spike counts than the lower spike counts seen in higher MNP output frequencies.

The results of the event-based CloudBrain simulation are consistent with the time-based NEST simulation showing that output frequency can be manipulated based on input to the NSI. This suggests that the NSI can translate both spiking and analog data into usable information for the network. The ability to receive spiking input increases the usability of the NSI as a possible encoding tool, indicating potential for interfacing with event-driven sensors. Additionally, the ability to run the NSI on an event-based simulation in real-time means that it can be applied in closed-loop control of robots.

## 5 Conclusion

The method introduced in our research is able to integrate NSI’s into SNNs to create a mixed network. Our model NSI is a biologically plausible input value encoder, receiving analog values as *I_input_* and passing the information to spiking neuron populations which naturally output spikes. This research confirms that the amplitude, frequency, and phase of an sCPG network can be manipulated based on changing input to an NSI, implying that an NSI can function as an encoding mechanism within an SNN. Furthermore, the ability of the network to adjust to an internal signal suggests that this setup could be useful in adaptive controllers.

Our recommended architecture for integrating an NSI with a mutually inhibitory sCPG network is shown in Figure 4, plot 4,5,6, allowing frequency to be adjusted by changing the voltage threshold potential. In this specific setup, the average frequency of the network can be regulated between 3.0*Hz* and 8.5*Hz*. Additionally, the regulation of synaptic conductance weight from the NSI allows an average peak output amplitude between approximately 6 and 52 spikes per time window. This assumes a rate-coding time window of 5*ms* and 5 neurons for the MNP as investigated in this study. The recorded frequency and amplitude ranges are determined by the architecture and parameter ranges so it would be advantageous to look further into the dynamics of this system to maximize flexibility. Additionally, the resolution of attainable frequencies should be evaluated as this will be an important metric for adaptive control.

An implementation of an NSI should be further researched to compare sub-threshold use of existing spiking neuron models to a unique non-spiking model. Further research into both the offset and injection current equations as well as their respective parameters is also necessary. Additionally, the leakage constant of the NSI could be optimized to ensure biological-plausibility and effectiveness. The relationships between frequency and phase as well as frequency and amplitude should be quantified so that these dependencies can be fully exploited.

The finding that amplitude can be scaled based on the value of the current injection to the MNP leads to the possibility of using an NSI to prioritize sensory inputs or inform coordination tasks. Insect inter-leg coordination is known to depend on sensory input not only from the local leg but also neighboring legs (Bidaye et al., 2018). Therefore, synaptic weights from a single NSI population could be tuned in order to inject more current to the local leg while also providing varying amounts of injection current to neighboring legs.

The event-based demonstration on the CloudBrain platform indicates that implementation on neuromorphic hardware is possible. A closed-loop mixed network should be further studied in CloudBrain since it can communicate with a robot in real-time (Larsen et al., 2020). This provides the opportunity for investigation of a mixed network adaptive controller using environmental interaction.

The ability of sub-threshold encoding to shape an sCPG network’s output opens up possibilities for new approaches to coordination and control tasks. Storchi et al. (2012) report observing the use of combinations of encoding mechanisms in biological systems, indicating that the use of multiple methods within a single robot could be a fruitful investigation direction for adaptive controllers.

## 6 Author Contributions

BS and LBL developed the main idea of the paper. BS researched the biological concepts, formulated a theory, and applied it within a time-based simulation tool. LBL and RKS conceived of the event-based NSI model and RKS implemented and validated neuron and synapse implementations. RKS implemented the sCPG in CloudBrain. The manuscript was written by BS with support from LBL and PM. LBL supervised the project and provided the funding.

