## Supplementary Material for "Integrating Non-Spiking Interneurons in Spiking Neural Networks"

Posted: August 14, 2020; Updated: November 26, 2020

bioRχiv
THE PREPRINT SERVER FOR BIOLOGY

### Integrating Non-Spiking Interneurons in Spiking Neural Networks

Beck Strohmer[1,*], Rasmus Karnøe Stagsted[1], Poramate Manoonpong[1], and Leon Bonde Larsen[1]

[1]SDU Biorobotics, Maersk McKinney Moller Institute, University of Southern Denmark

#### Supplementary Material

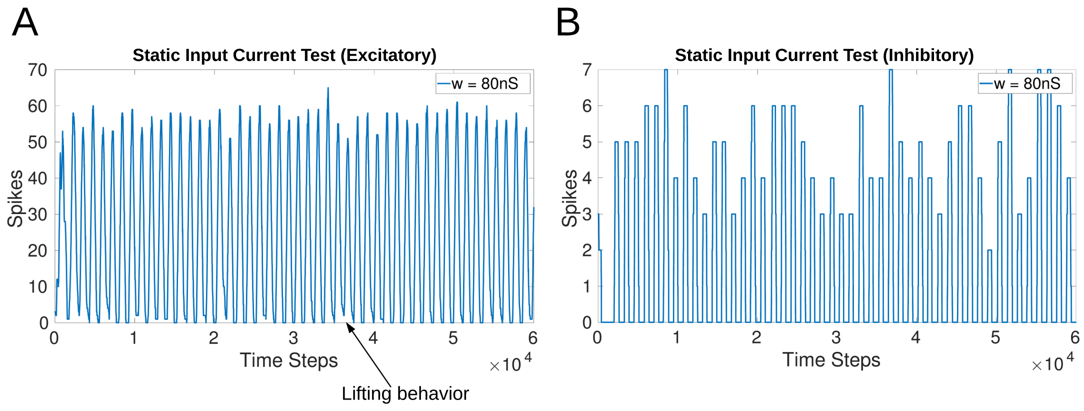


Figure 1: A and B: Results for a conductance weight of $w = 80nS$ and a constant input current to the NSI of $148pA$. A lifting behavior is noticeable within some time windows of the excitatory test (A) where the rate-coded output does not return to zero. The inhibitory test (B) output is initially suppressed.

Table 1: Start and end frequency per voltage characteristic manipulation test, only tests involving both sCPG neuron populations are shown. Tests for $V_{th}$ and $V_{reset}$ show frequency can be increased by a factor of 3. The $V_m$ tests are inconclusive because they are unstable.

| Frequency Comparison | | |
|---|---|---|
| Test | Start Frequency (Hz) | End Frequency (Hz) |
| 4_0 ($V_{th}$ exc) | 2.77 | 8.47 |
| 4_3 ($V_{th}$ inh) | 2.36 | 8.01 |
| 5_0 ($V_{reset}$ exc) | 2.94 | 8.94 |
| 5_3 ($V_{reset}$ inh) | 2.45 | 8.15 |
| 6_0 ($V_m$ exc) | 8.14 | 5.32 |
| 6_3 ($V_m$ inh) | 0.89 | 8.19 |

Table 2: Average maximum number of spikes based on the MNP output frequency. A static input current is provided to the NSI which changes the voltage threshold of the sCPG populations through an offset leading to different output frequencies.

| Average Maximum Spikes based on MNP Output Frequency | | | |
|---|---|---|---|
| $I_{input}$ (pA) | $V_{th}$ (mV) | Average Freq (Hz) | Average Max Spikes |
| 0.0 | -55.6 | 3.0 | 96.6 |
| 29.6 | -54.7 | 4.1 | 89.7 |
| 59.2 | -53.7 | 5.2 | 70.0 |
| 88.8 | -52.7 | 6.5 | 54.3 |
| 118.4 | -51.7 | 7.5 | 51.5 |
| 148.0 | -50.7 | 8.5 | 49.1 |

In all of the following plots, excitatory and inhibitory tests are compared to their respective counterparts. The vertical lines indicate when a change in input current occurs. The exact configuration details for each test are found in Tables 1 and 2 and visualized in Figures 4 and 5.

#### Tests Involving Both sCPG Neuron Populations

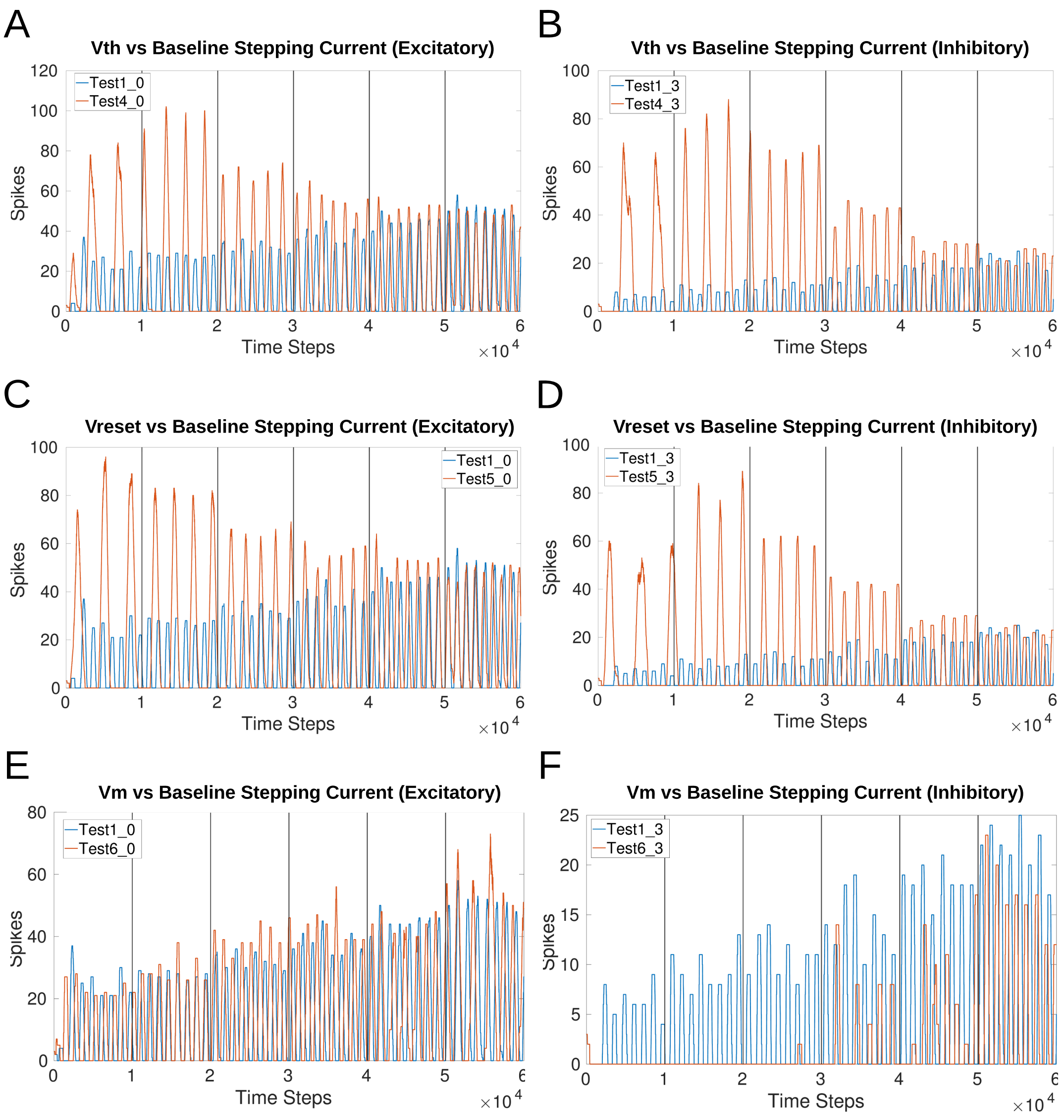


Figure 2: Comparison of voltage characteristic manipulation to the baseline MNP current injection step test. Outputs show that frequency is linearly affected by updating $V_{th}$ and $V_{reset}$ of sCPG neuron populations. Adding an offset to $V_m$ of the post-synaptic neuron populations creates instability, the frequency of the signal does not increase linearly in comparison to the change in membrane potential. A and B: Voltage threshold is updated and current is injected to the MNP; C and D: Voltage reset value is updated and current is injected to the MNP; E:and F: Membrane potential is updated and current is injected to the MNP;

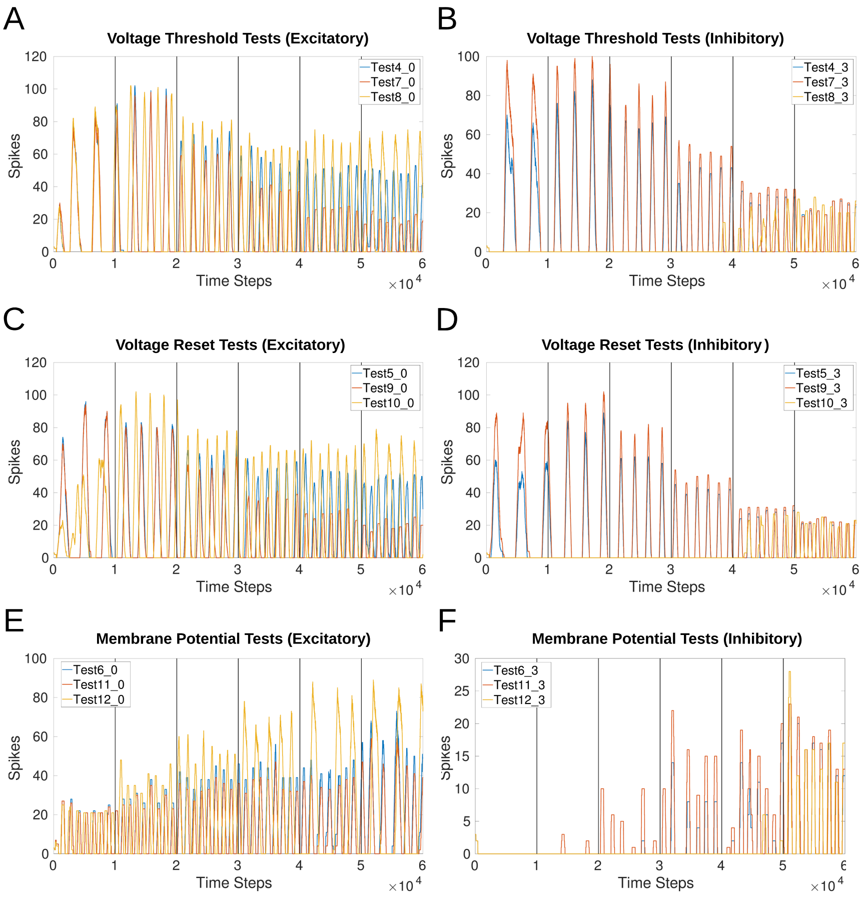


Figure 3: Comparison of voltage characteristic manipulation to the sCPG neuron populations with and without current injection to the sCPG neuron populations and MNP. Network behavior remains the same when current is injected to the MNP during frequency changes for all voltage characteristic manipulations. Current injection to the sCPG populations adds a phase shift and changes the output frequency. A and B: Voltage threshold is updated, Test 4 injects current to the MNP, Test 7 does not inject current to any post-synaptic populations, and Test 8 injects current to both the sCPG populations and the MNP; C and D: Voltage reset value is updated, Test 5 injects current to the MNP, Test 9 does not inject current to any post-synaptic populations, and Test 10 injects current to both the sCPG populations and the MNP; E and F: Membrane potential is updated, Test 6 injects current to the MNP, Test 11 does not inject current to any post-synaptic populations, and Test 12 injects current to both the sCPG populations and the MNP;

#### Tests Involving Single sCPG Neuron Populations

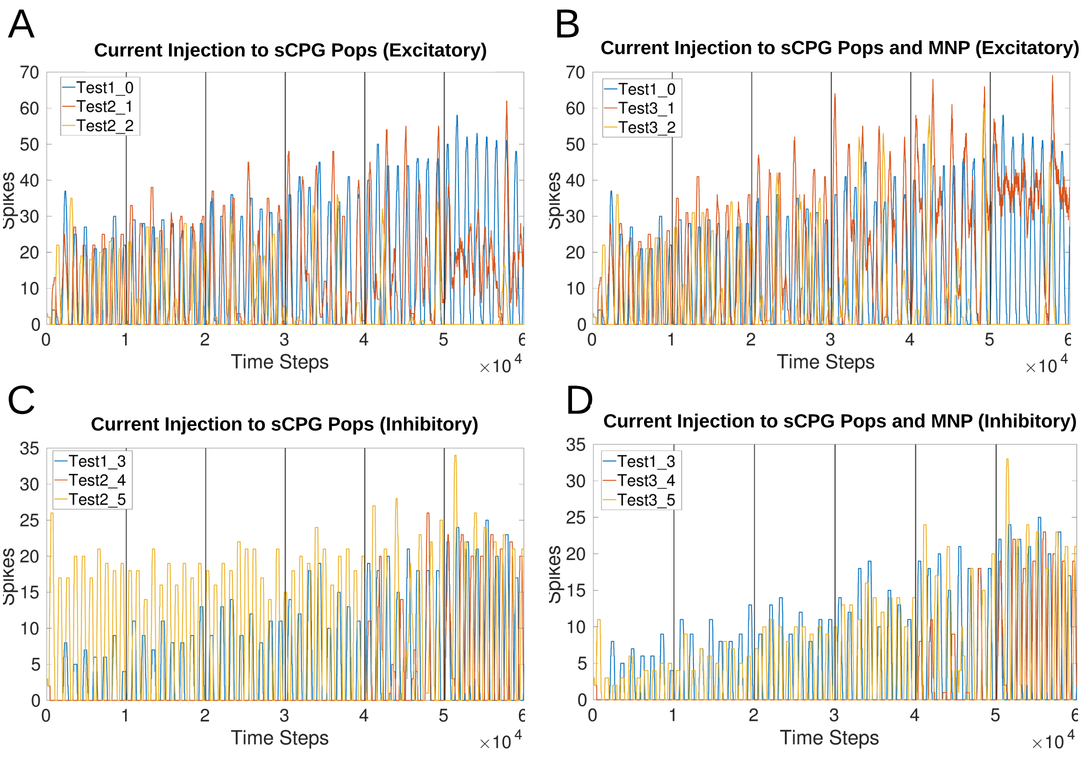


Figure 4: Comparison of current injections to either the excitatory (_1 and _4) or inhibitory (_2 and _5) sCPG neuron population to the baseline MNP current injection step test. Outputs show that current injection to a single sCPG population creates instability by over-inhibition suppressing output or over-excitation resulting in the signal not returning to zero. A and C: Current injection to the sCPG populations only.; B and D: Current injection to the sCPG populations and the MNP;

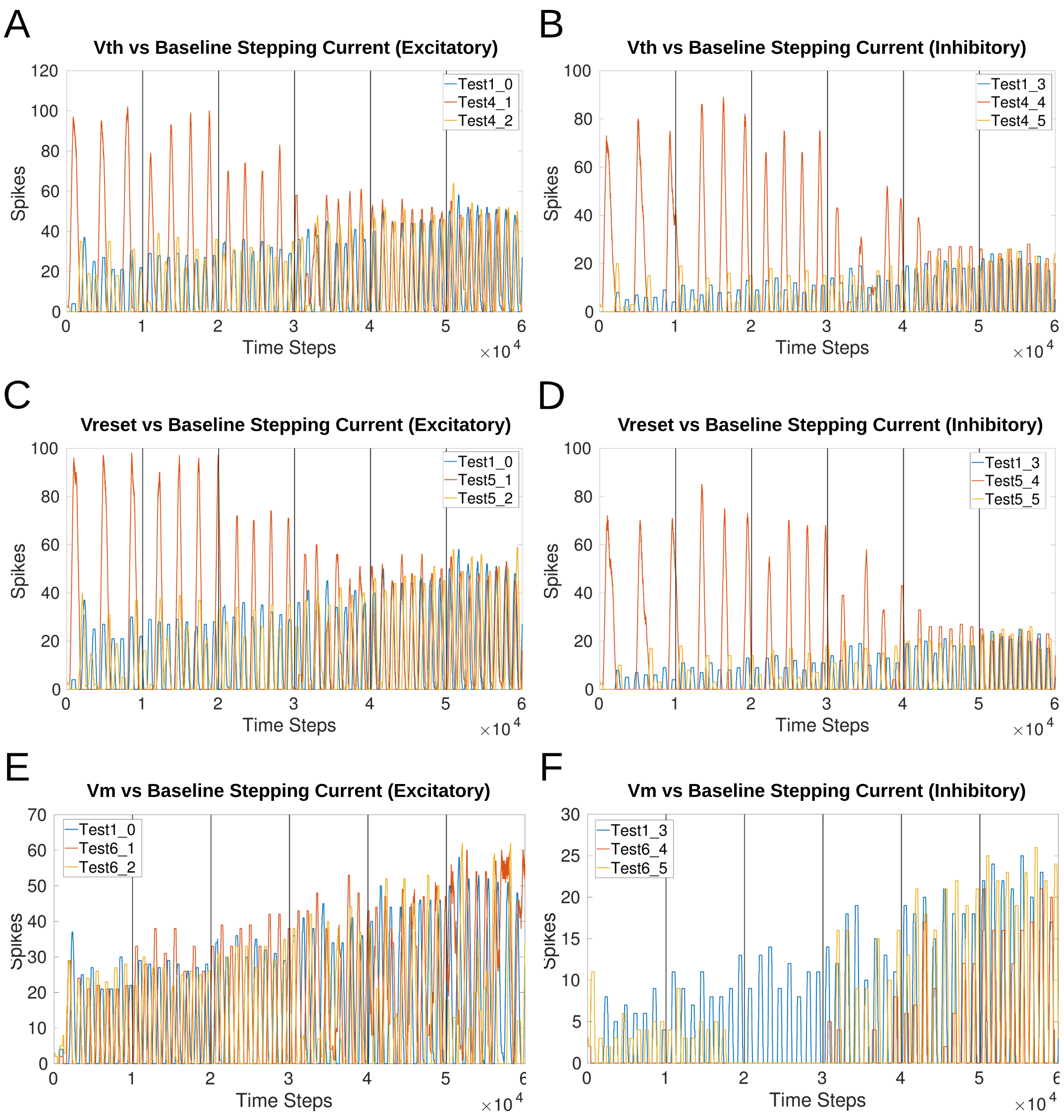


Figure 5: Comparison of voltage characteristic manipulation of either the excitatory (_1 and _4) or inhibitory (_2 and _5) sCPG neuron population to the baseline MNP current injection step test. Frequency is affected by updating $V_{th}$ and $V_{reset}$ of a single sCPG population but there are occasional "double peaks". Adding an offset to $V_m$ of a single post-synaptic population changes the output characteristics of the signal. A and B: Voltage threshold is updated and current is injected to the MNP; C and D: Voltage reset value is updated and current is injected to the MNP; E and F: Membrane potential is updated and current is injected to the MNP;

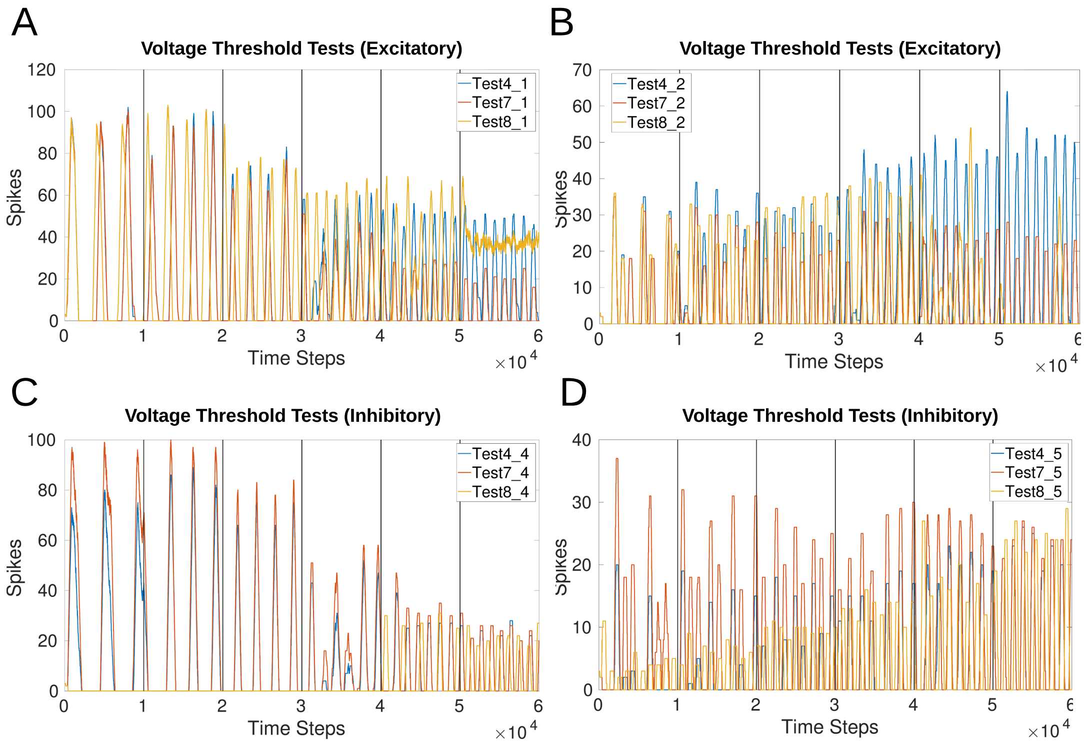


Figure 6: Comparison of $V_{th}$ manipulation to either the excitatory (_1 and _4) or inhibitory (_2 and _5) sCPG neuron population. Test 4 injects current to the MNP, Test 7 does not inject current to any post-synaptic populations, and Test 8 injects current to both the sCPG populations and the MNP. A and B: Tests with excitatory current injection from the NSI; C and D: Tests with inhibitory current injection from the NSI;

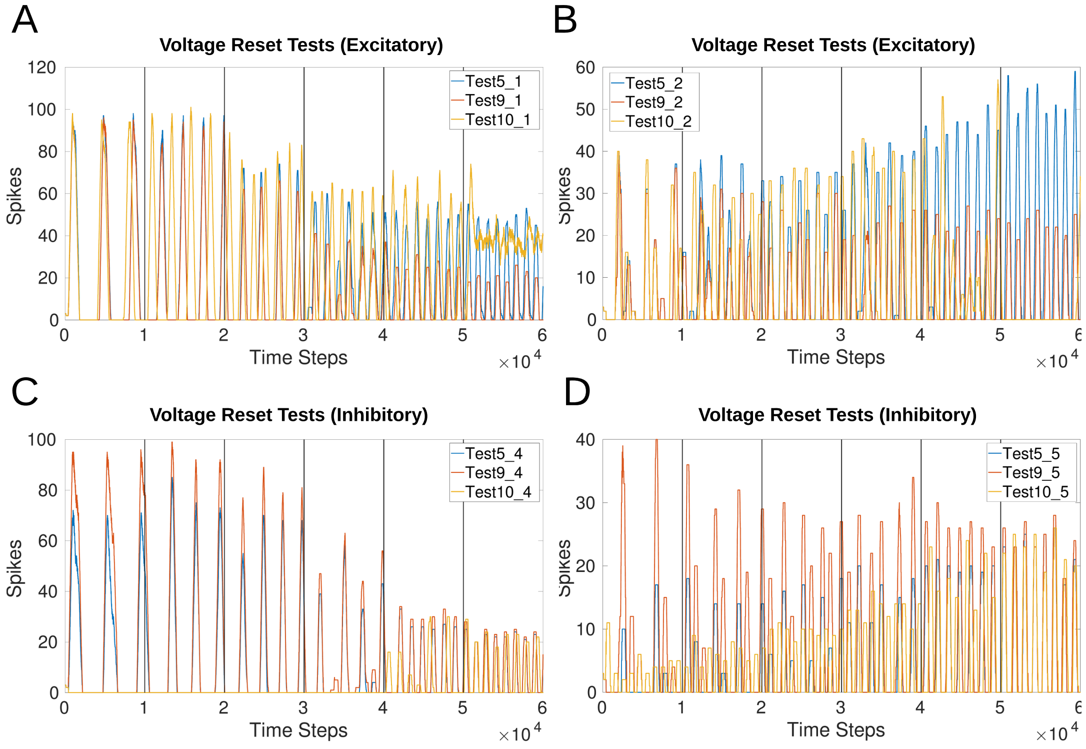


Figure 7: Comparison of $V_{reset}$ manipulation to either the excitatory (_1 and _4) or inhibitory (_2 and _5) sCPG neuron population. Test 5 injects current to the MNP, Test 9 does not inject current to any post-synaptic populations, and Test 10 injects current to both the sCPG populations and the MNP. A and B: Tests with excitatory current injection from the NSI; C and D: Tests with inhibitory current injection from the NSI;

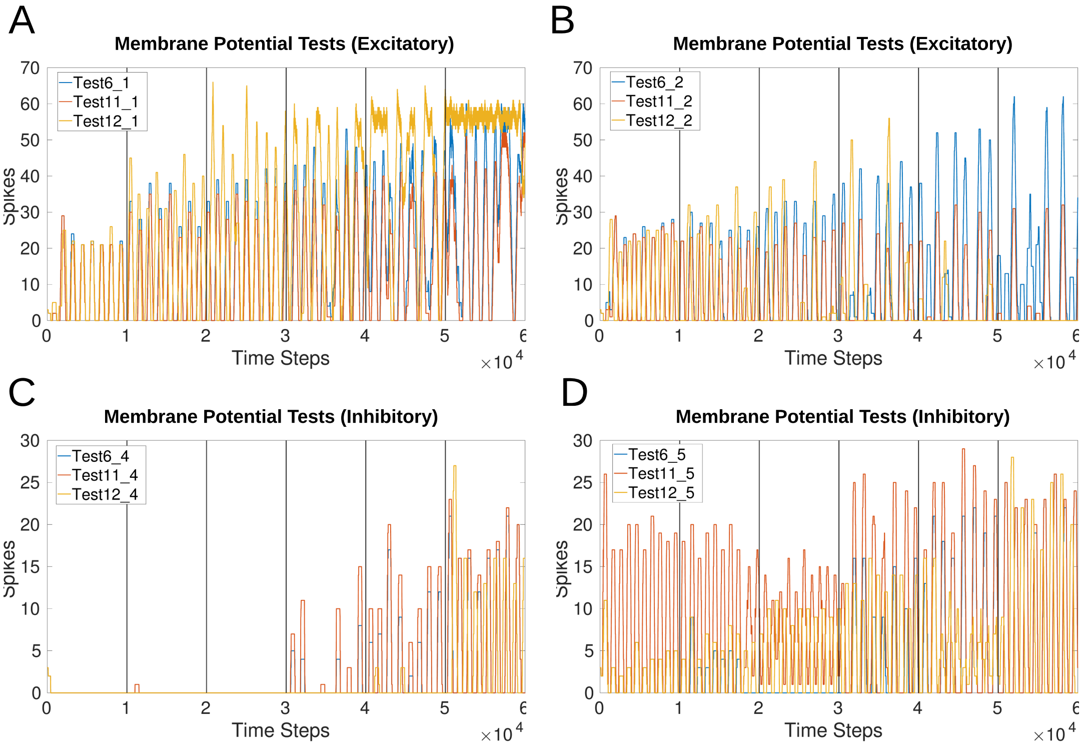


Figure 8: Comparison of $V_m$ manipulation to either the excitatory (_1 and _4) or inhibitory (_2 and _5) sCPG neuron population. Test 6 injects current to the MNP, Test 11 does not inject current to any post-synaptic populations, and Test 12 injects current to both the sCPG populations and the MNP. A and B: Tests with excitatory current injection from the NSI; C and D: Tests with inhibitory current injection from the NSI;
